# Force-induced site-specific enzymatic cleavage probes reveal that serial mechanical engagement boosts T cell activation

**DOI:** 10.1101/2023.08.07.552310

**Authors:** Jhordan Rogers, Rong Ma, Yuesong Hu, Khalid Salaita

## Abstract

The surface of T cells is studded with T cell receptors (TCRs) that are used to scan target cells to identify peptide-major histocompatibility complexes (pMHCs) signatures of viral infection or cancerous mutation. It is now established that the TCR-pMHC complex is highly transient and experiences mechanical forces that augment the fidelity of T cell activation. An important question in this area pertains to the role of force duration in immune activation. Herein, we report the development of force probes that autonomously terminate tension within a time window following mechanical triggering. Force-induced site-specific enzymatic cleavage (FUSE) probes tune tension duration by controlling the rate of a force-triggered endonuclease hydrolysis reaction. This new capability provides a method to study how accumulated force duration contributes to T cell activation. We screened DNA sequences and identified FUSE probes that disrupt mechanical interactions with *F* >7.1 piconewtons (pN) between TCRs and pMHCs. Force lifetimes (τ_F_) are tunable from tens of min down to 1.9 min. T cells challenged with FUSE probes presenting cognate antigens with τ_F_ of 1.9 min demonstrated dampened markers of early activation, thus demonstrating that repeated mechanical sampling boosts TCR activation. Repeated mechanical sampling *F* >7.1 pN was found to be particularly critical at lower pMHC antigen densities, wherein the T cell activation declined by 23% with τ_F_ of 1.9 min. FUSE probes with *F* >17.0 pN response showed weaker influence on T cell triggering further showing that TCR-pMHC with *F* >17.0 pN are less frequent compared to *F* >7.1 pN. Taken together, FUSE probes allow a new strategy to investigate the role of force dynamics in mechanotransduction broadly and specifically suggest a model of serial mechanical engagement in antigen recognition.

## Introduction

Cytotoxic, or CD8+, T cells are essential during the adaptive immune response, as they are responsible for identifying and eradicating virally infected or cancerous cells. The T cell receptor (TCR) distinguishes between non-stimulatory ‘self’ peptide and stimulatory ‘foreign’ peptide antigens that are presented by major histocompatibility complexes (MHCs) on the surface of virtually every cell type. This discrimination process is extremely effective, even though there is minimal difference between the affinities of the non-stimulatory and stimulatory peptide MHCs (pMHCs), as both bind to the TCR with a K_D_ typically in the μM regime, which is among the weakest receptor-ligand interactions in biology.^1-6^ Additionally, T cells are ultrasensitive; CD8+ T cells can become activated by as few as one to three stimulatory pMHCs on the surface of an antigen presenting cell.^7-10^ Although T cells are highly sensitive and specific towards aberrant cells, the molecular mechanisms that initiate their cytotoxic effector functions remain poorly understood.

To help explain the phenomenal specificity of the TCR, one prominent model suggests that the TCR functions as a mechanosensor; mechanical forces transmitted to the TCR-pMHC complex boosts its discriminatory power.^11-15^ Early single-molecule experiments showed enhanced T cell activation in response to the application of 10 pN force applied to the TCR-pMHC. These small, fine-tuned forces on the scale of 5-20 pN have also been suggested to stabilize the interaction between the TCR and pMHC, ultimately increasing the lifetime of the bond.^16-19^ Our group provided evidence validating the mechanosensor model by developing sensors that mapped ∼12-19 pN T cell forces generated by the T cell cytoskeleton and transmitted to their TCR-antigen bonds during antigen recognition.^20-22^ Briefly, molecular force probes visualize TCR forces by presenting pMHC ligands conjugated to fluorescently labeled DNA hairpins that are immobilized onto a glass coverslip (**Figure S1**). These probes contain a fluorophore-quencher pair attached to the termini of the hairpin stem. Once a TCR binds to the antigen and exerts *F* greater than the F_1/2_ of the hairpin (50% probability of unfolding the hairpin at equilibrium), then the fluorophore is separated from the quencher, leading to a 100-fold increase in fluorescence intensity (**Figure 1A**).

**Figure 1.**
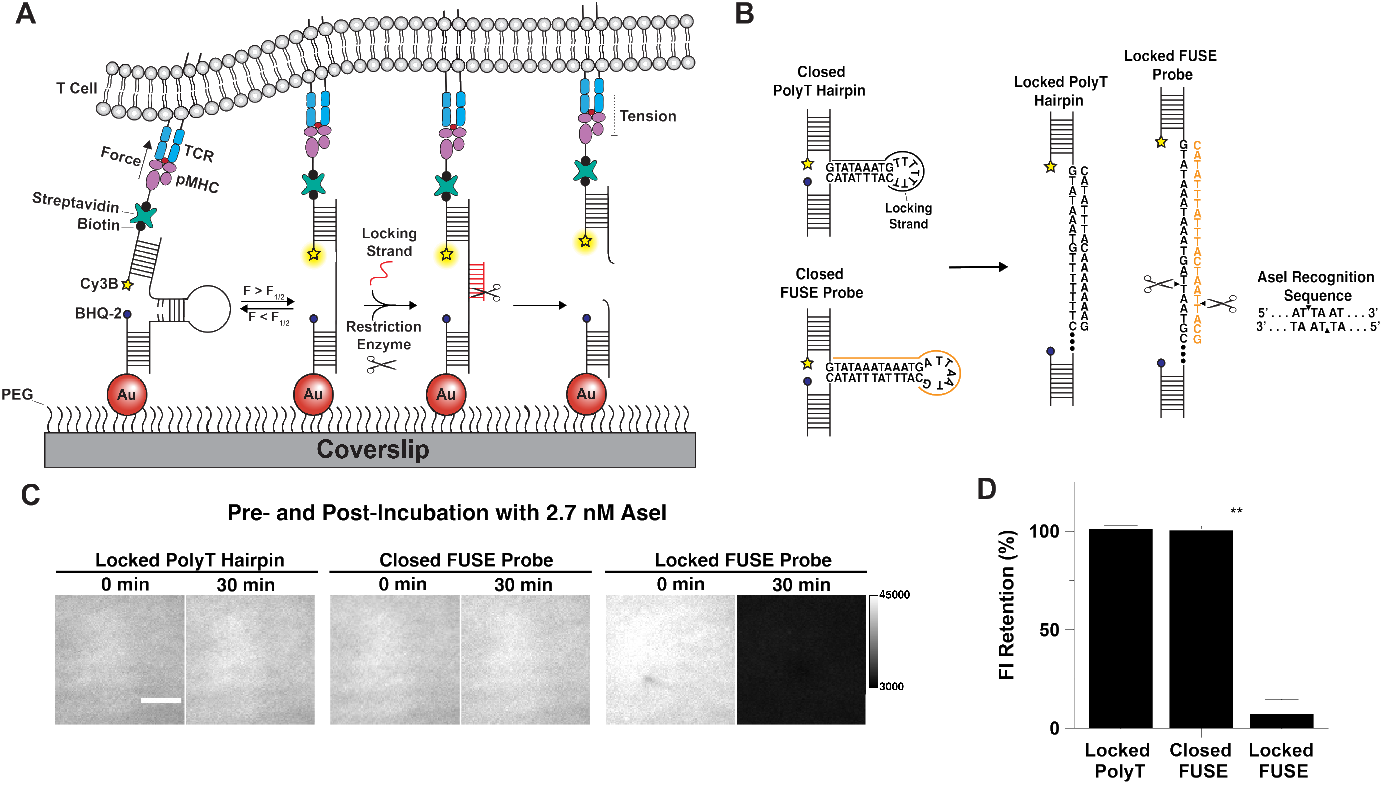
A) Schematic of force-induced site-specific enzymatic cleavage (FUSE) assay. B) Design of the FUSE probe compared to the traditional 4.7 pN probe with a polyT loop. The 21mer locking strand for the FUSE probe (orange) binds to the open hairpin to complete the recognition site for AseI. C) Representative TIRF images of the locked 4.7 pN probe with polyT loop, closed FUSE probe, and locked FUSE probe before and after adding AseI restriction endonuclease. Scale bar = 5 μm. D) Quantification of the retention of fluorescence intensity with surfaces presenting either the locked 4.7 pN probe with polyT loop (102±1%), the closed FUSE probe (101±1%), or the locked FUSE probe (8±7%) with 2.7 nM AseI. Statistical analysis performed using Student’s t test, **P<0.01.

Another prominent model that helps explain TCR sensitivity is the serial engagement model which postulates that TCRs repeatedly engage stimulatory pMHCs to trigger activation at low antigen density.^23-28^ Kalergis et al. have demonstrated a “Goldilocks-like” relationship between TCR-pMHC dwell time and T cell activation. If dwell times are too short, then the TCRs fail to initiate signaling. However, if dwell times are too long, then few TCRs benefit from serial engagement and T cell activation is dampened.^3^ TCR clustering may also further promote serial engagement through rapid rebinding, as it has been suggested that pMHCs interacting with TCR microclusters can serially bind multiple TCRs before diffusing away along the cell membrane.^29, 30^ Despite the accumulating experimental support for both the serial engagement model and the mechanosensor model, it remains unclear how these models work together.

Accordingly, the primary goal of this work is to develop a tool to explore how these two models may operate together to enhance TCR triggering. Here, we introduce force-induced site-specific enzymatic cleavage (FUSE) probes, which allows one to tune the duration of TCR-pMHC force (**Figure 1A**). FUSE probes contain the core components of DNA hairpin molecular force probes but are degraded after the antigen is mechanically sampled. This process occurs by adding a single-stranded DNA (locking strand) that is complementary to the cryptic loop region of the hairpin that is only exposed once the probe is unfolded.^22^ This cryptic region contains a sequence that is recognized by a site-specific restriction endonuclease only when the locking strand hybridizes to the unfolded probe. This selective cleavage disrupts the mechanical resistance of the pMHC, as it is no longer tethered to the surface by a DNA duplex. The rate of this disruption is tuned by varying the concentration of nuclease added, which allows the duration of mechanical resistance, or the tension duration experienced by TCRs, to be controlled orthogonally and without modification to the antigen or TCR.

We demonstrate that the cleavage of FUSE probes with the AseI restriction endonuclease is highly site-specific. Surface rate measurements of the locking strand binding to FUSE probes, as well as the rate of enzymatic cleavage of the locked probe showed that FUSE probe cleave is highly selective, and closed probes are cleaved at a rate 10^4^ lower than that of opened probes. Force lifetimes following triggered are tunable down to 1.9 min which was achieved at 10.7 nM AseI. FUSE probes presenting a pMHC loaded with the ovalbumin-derived peptide, SIINFEKL, were used to demonstrate that FUSE probe cleavage was unperturbed at the junction between a T cell and antigen-coated substrate. Critically, antigen-FUSE probes terminate tension but remain confined at the T-cell surface because of rebinding to high density TCRs as we measured a half-life of 12.8 min. Lastly, early T cell signaling, as reflected by the ZAP70 phosphorylation level at *t*=15 min, is dampened with 7.1 pN FUSE probes, and this dampening is further enhanced at low antigen density. These results lead to the conclusion that serial mechanical engagement boosts T cell activation.

## Results

### Characterization of force-induced site-specific enzymatic cleavage (FUSE) probes

FUSE probes differ from traditional hairpin probes in their force-triggered self-cleavage response. We aimed to achieve this function by incorporating a recognition sequence for a site-specific restriction endonuclease in the loop segment of the DNA hairpin that can only be cleaved upon mechanical melting of the hairpin (**Figure 1B**). Specifically, the endonuclease recognition sequence is exposed only once a complementary ‘locking strand’ hybridizes to the mechanically unfolded probe. Thus, the locking strand must exhibit two features: the first is rapid and specific hybridization to unfolded FUSE probes. The second is that the ‘locked probe’ must demonstrate high thermostability at 37°C for optimal enzymatic activity (**Figure S2A**). To meet these criteria, we designed and screened a small library of FUSE sequences (**Table S1** and **Figure S2B**).^31, 32^ Our initial sequences were unsuccessful as we found that FUSE probes incorporating our previously reported 4.7 pN probe design (stem = 22% GC, 9 bp) resulted in poor thermostability at 37°C once it was locked with its accompanying 15-nt locking strand (**Figure S2C**).^22^ Ultimately, we found that by extending the stem of the hairpin to 13 bp, replacing the 7^th^ T base in the loop with a G, and increasing the locking strand length to 21-nt, led to increasing the stability of the locked probe substantially. The optimized sequence is shown in **Figure 1B**, and the F_1/2_ of this sequence was 7.1 pN as calculated using established precedent (**Equation S1** and **S2**).^20, 33, 34^ Since site-specific enzymatic cleavage is imperative for specifically perturbing mechanical interactions, we next aimed to quantify AseI activity and specificity. Here, we introduced AseI to a surface tethered locked 4.7 pN (non-specific) probe with a polyT loop, closed 7.1 pN FUSE probe, and locked FUSE probe.^21, 22^ Note that the locked probes were labeled with a Cy3B-BHQ-2 fluorophore quencher pair, while the closed 7.1 pN FUSE probe only included the Cy3B reporter to facilitate quantifying cleavage rates. By measuring the time-dependent decay of the Cy3B signal associated with the top strand of each probe, we were able to determine the specificity and activity of AseI on surface tethered substrate. Importantly, under our conditions, there was no observable cleavage of the locked non-specific hairpin probe with a polyT loop that was lacking the recognition sequence. The closed FUSE probe, which only contains the incomplete recognition sequence also did not show any detectable cleavage (**Figure 1C**). In contrast, 90% of the locked FUSE hairpin was cleaved under identical conditions after a 30 min incubation at 37°C with 2.7 nM AseI (**Figure 1C** and **1D**).

Next, we aimed to quantify the kinetics of FUSE triggering. This includes nonspecific hybridization between a closed FUSE probe and its locking strand, specific hybridization between an unfolded FUSE hairpin and its locking strand, and the rate of enzymatic cleavage of the locked probe (**Figure 2** and **S3**). To determine the lock hybridization kinetics, we coated coverslips with either the closed FUSE probe or the ‘unstructured hairpin’, which is a single-stranded DNA that only contains the portion of the hairpin that is complementary to the locking strand. We then added either unlabeled or fluorophore-labeled locking strand and observed the increase in the signal associated with the hairpin opening or locking strand binding to unstructured hairpin (**Figure 2A** and **2D**). The increase in this signal can be fitted to a one phase association curve (**Equation S3**), which allows the apparent rate, k, to be determined for a given concentration of locking strand (**Figure 2B, 2E, S2B**, and **S2D**). We then plotted the rate versus locking strand concentration, which allowed us to extrapolate a rate for any given locking strand concentration, assuming pseudo-first order kinetics (**Figure 2C** and **2F**). To quantify the rate of enzymatic cleavage of the locked probe, the FUSE probe was annealed with its locking strand prior to conjugation to the surface (**Figure 2G**). Then, AseI restriction endonuclease concentrations ranging from 0.5 nM–10.7 nM were incubated at 37°C for 30 min (**Figure S3F**). The enzymatic activity of AseI was observed by tracking the decrease in signal as the fluorophore-labeled top strand diffuses away from the surface after cleavage. This decay was fit to a one phase exponential decay curve (**Equation S4**) to determine the half-life (t_1/2_) and average lifetime of the locked probe (τ_F_) (**Figure 2H** and **S3F**). We then plotted this rate of decay versus AseI concentration to compare enzymatic activity across a range of endonuclease concentrations (**Figure 2I**). For FUSE probes to terminate mechanical signaling accurately and rapidly, the specific hybridization and enzymatic cleavage rates must be much faster than the rates of nonspecific hybridization and cleavage.

**Figure 2.**
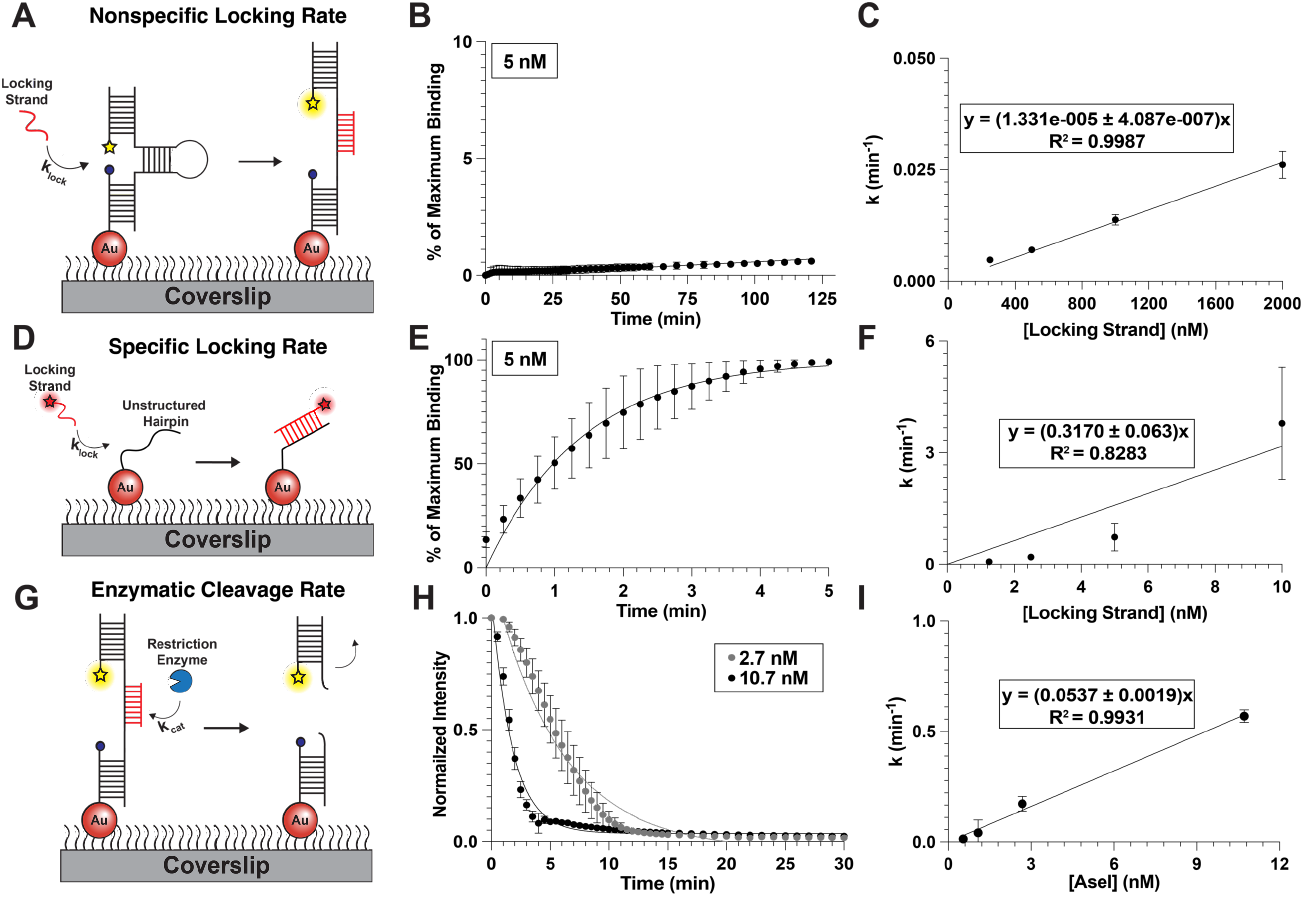
A) Schematic of experiment to quantify rate of nonspecific hybridization of locking strand. B) Representative plot of the percentage of maximum locking strand binding to the hairpin versus time to quantify the nonspecific rate of hybridization for 5 nM of locking strand, k = 5.75e-005 ± 3.4e-006 min^-1^. C) Plot of the pseudo first-order rate constant of nonspecific hybridization versus concentration of locking strand. D) Schematic of experimental design to quantify the rate of specific hybridization of locking strand to its exposed docking site. E) Representative plot of the percentage of maximum locking strand binding to the unstructured hairpin versus time to quantify the specific rate of hybridization for 5 nM of locking strand, k = 0.730 ± 0.374 min^-1^. F) Plot of the pseudo firstorder rate constant for specific hybridization versus the concentration of locking strand. G) Schematic displaying the experimental setup to quantify the rate of enzymatic cleavage of locked hairpin. H) Representative plot of the normalized surface intensity versus time to determine the rate of cleavage and τ_F_ for 2.7 and 10.7 nM of AseI. The apparent rate of cleavage for 2.7 nM AseI was 0.173 ± 0.0.034 min^-1^ and 0.570 ± 0.0.029 min^-1^ for 10.7 nM. I) Plot of the apparent rate of cleavage versus concentration of AseI.

Indeed, we did observe a 2.4×10^4^ faster specific hybridization rate (1585 ± 315 min^-1^) compared to the nonspecific hybridization rate (0.066 ± 0.006 min^-1^) for 5 μM of locking strand, which is the concentration we chose for the FUSE assay (**Figure 2B, 2C, 2E**, and **2F**). Additionally, we found that the τ_F_ of the locked probe was 5.8 ± 0.7 min at 2.7 nM and 1.9 ± 0.3 min at 10.7 nM of AseI. These values suggest that FUSE probes are able to be effectively locked and cleaved upon mechanical triggering.

### FUSE probes can dynamically map and disrupt TCR-ligand mechanical interactions

After validating the kinetic parameters of these probes with cell-free experiments, we next wanted to ensure FUSE probes can be employed to study receptor-ligand interactions. Although the prior cell-free experiments demonstrated the feasibility of this assay, it was unclear whether the cellular environment would impede the enzyme’s ability to access the locked substrate. For these experiments, we elected to use the well-studied OT-1 transgenic T cell model, where TCRs are reactive to the ovalbumin SIINFEKL peptide (N4). We decorated 7.1 pN FUSE probes with N4 pMHC ligand.^1, 16, 21, 22, 35^ The F_1/2_ of 7.1 pN was chosen because this force magnitude is well within the range that OT-1 TCRs transmit to the N4 pMHCs.^16, 22^ Briefly, we allowed the OT-1 cells to spread on coverslips for ∼10-15 min with either fluorophore-labeled or unlabeled locking strand to allow locked tension signal to accumulate (**Figure S4**). We then added or withheld AseI and tracked the signal of either the fluorophore associated with the locking strand (Atto647N), or the fluorophore associated with the top strand/pMHC (Cy3B) underneath cells (**Figure 3A** and **3B**). In **Figure 3B**, both the Atto647N and Cy3B signals are associated with tension; however, the decay of the Atto647N signal reports the rate of enzymatic cleavage of the locked probe. Since the Cy3B dye is conjugated to the antigen strand, its decay is hindered due to rebinding to TCRs that may occur after the probe’ is released from the surface. Indeed, the rate of decay of the lock signal (Atto647N) is robust with only 15% of the initial signal remaining under cells after AseI is added (**Figure 3C**). While this depletion is apparent, this decay happens at a slightly slower rate compared to the cell-free cleavage assay (τ = 6.9 min) (**Figure 3D**). In contrast, the fluorophore associated with the ligand (Cy3B) depletes at a much slower rate (τ= 18.5 min) and around 30% of the initial signal remains after a 20 min incubation with AseI (**Figure 3E** and **3F**). We speculate that this slower rate of Cy3B depletion is due to TCRs rebinding pMHCs once these probes are cleaved from the surface. Altogether, these results show that FUSE probes are effectively cleaved at the T cell-substrate junction and demonstrate extended dwell time at the T cell surface despite termination of mechanical resistance.

**Figure 3.**
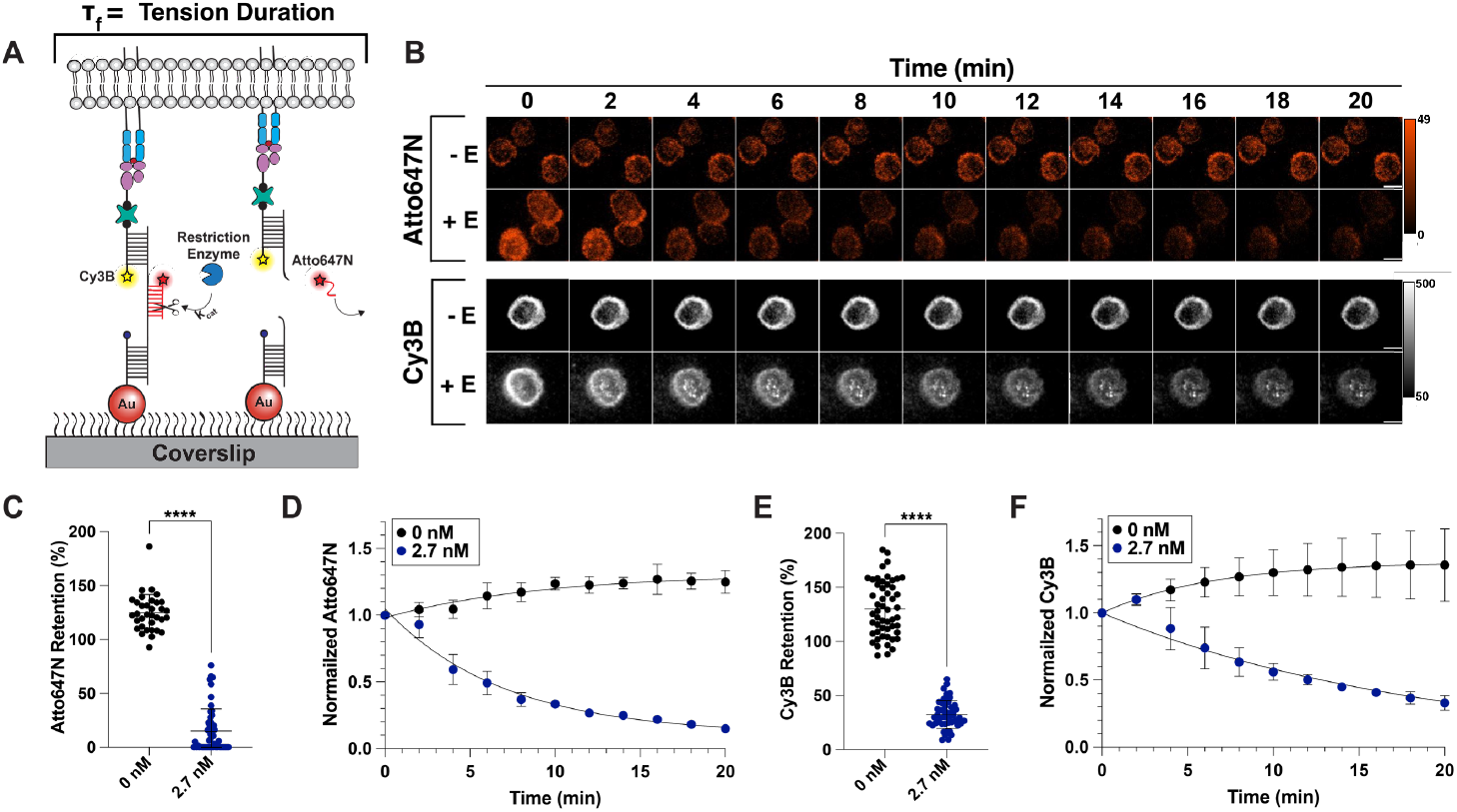
A) Schematic showing the cleavage of locked FUSE probes under cells; the rate of this enzymatic cleavage defines the average tension duration experience by TCRs. B) Representative timelapse showing the decay of locked FUSE probes under cells. Atto647N signal tracks the locking strand, while Cy3B signal is used to visualize the ligand and top strand of the FUSE probe. Scale bar = 5 μm. C) Quantification of the change in locking strand signal under cells after incubation with or without AseI enzyme, n>30 cells for each condition. Statistical analysis was performed using Student’s t test, ****P<0.0001. D) Plot showing the change in lock signal under cells over the course of a 20-minute incubation with (blue) or without (black) AseI. E) Quantification of the depletion of ligand under cells after incubation with or without AseI, n>30 cells for each condition. Statistical analysis was performed using Student’s t test, ****P<0.0001. F) Plot showing the exponential decay of ligand under cells over the span of 20 min in the presence of AseI (blue) and a slight increase in tension signal in the absence of AseI (black).

### TCR-pMHC tension duration influences T cell activation

After demonstrating that FUSE probes are triggered and selectively cleaved in response to *F* ≥7.1 pN, we aimed to test the role of accumulated TCR-pMHC tension duration on T cell signaling.

Specifically, we were interested in quantifying early T cell activation in response to TCR-pMHC tension duration. To achieve this, we measured phosphorylation of the proximal kinase ZAP70 (pYZAP70) in OT-1 cells interacting with N4 antigen presented by FUSE probes. By titrating enzyme concentration, we were able to vary the average length of time that TCRs can mechanically interact with surface bound pMHCs, otherwise known as the τ_F_ (**Figure 4A)**. The maximum τ_F_ is achieved when no enzyme is added, as the TCR can physically engage with surface bound pMHCs until the cells are eventually fixed and stained to quantify intracellular pYZAP70. We used this level of activation as a control to compare pYZAP70 expression in conditions where either 10.7 nM (τ_F_ = 1.9 min) or 2.7 nM (τ_F_ = 5.8 min) of AseI was added with cells along with 5 μM of locking strand. Interestingly, we observed a 15% decrease in pYZAP70 expression when the τ_F_ was reduced to 1.9 min, but only an 8% decrease when τ_F_ was reduced to 5.8 min (**Figure 4B** and **4C**). Additionally, we limited the τ_F_ of FUSE probes presenting antiCD3ε to 5.8 min and observed a more pronounced reduction in pYZAP70 (15%) compared to probes presenting pMHC with the same τ_F_ (8%) (**Figure S5**). We hypothesize that this larger decrease in the antiCD3ε condition was due to the disruption of long-lived mechanical interactions between antiCD3ε and the TCR, as antiCD3ε displays a higher affinity (nM K_D_) towards the TCR than pMHCs (μM K_D_).^36, 37^ Note that both the AseI hairpin and accompanying locking strand must be present to observe a statistically significant decrease in T cell signaling, thus validating that selective FUSE probe cleavage is responsible for this perturbation (**Figure S6**).

**Figure 4.**
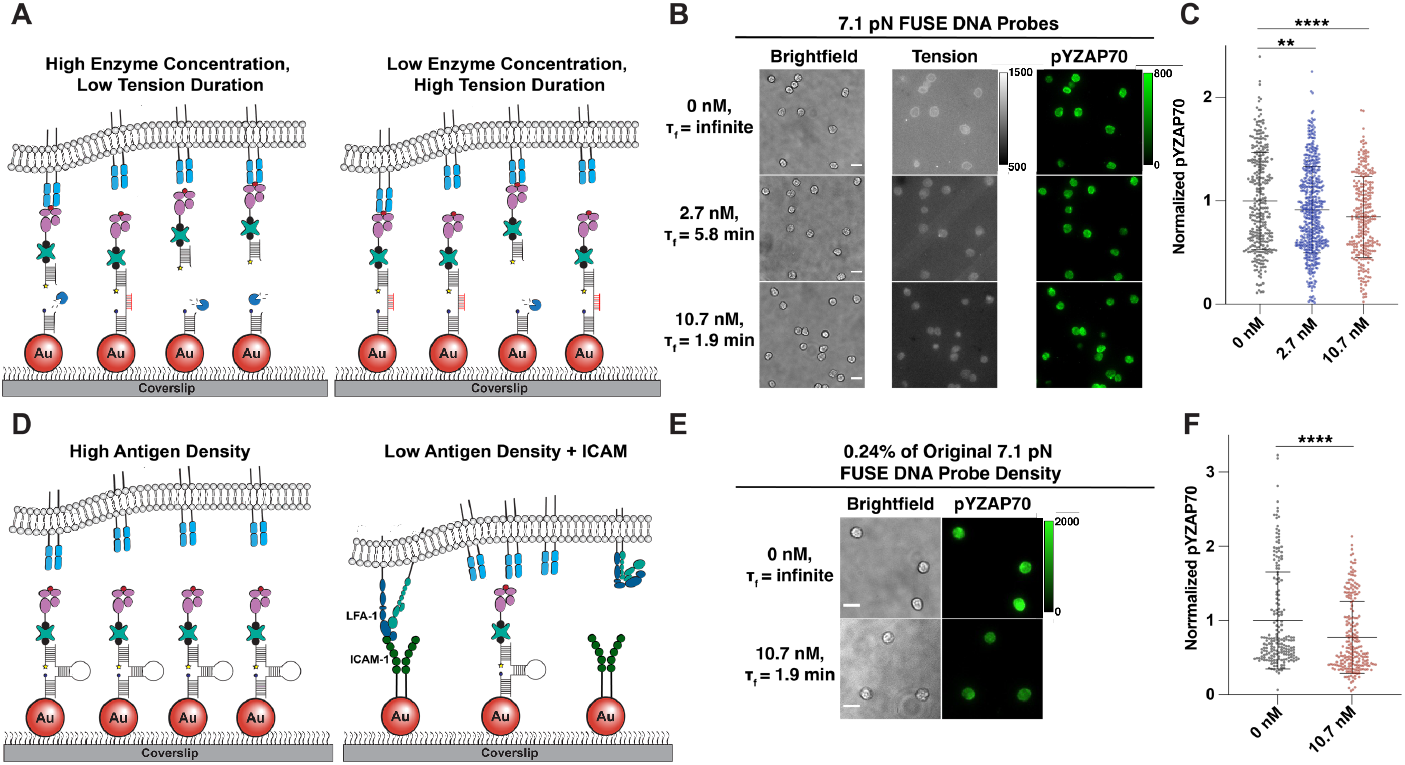
A) Schematic depicting the difference in TCR-pMHC interactions when presented on FUSE probes in high and low enzyme conditions. B) Representative images showing the difference in tension and pYZAP70 signal after a 15-minute FUSE assay with three different concentrations of enzyme (0 nM, 2.7 nM, 10.7 nM). Scale bar = 10 μm. C) Quantification of pYZAP70 signal after OT-1 cells were incubated with 0 nM (Mean norm. pY = 1.00), 2.7 nM (mean norm. pY = 0.92), and 10.7 nM (mean norm. pY = 0.85) of AseI on FUSE probes, n>300 cells. Statistical analysis was performed using Student’s t test between no enzyme control and experimental groups, **P<0.01 and ****P<0.0001. D) Schematic depicting cell interactions on a high density versus a low antigen density surface. E) Representative images showing the difference in pYZAP70 signal after a 15-minute FUSE assay in the presence and absence of 10.7 nM AseI. Scale bar = 10 μm. F) Quantification of pYZAP70 signal in cells after incubation with 10.7 nM AseI (mean norm. pY = 0.77) and without AseI (mean norm. pY = 1.00), n>200 cells. Statistical analysis was performed using Student’s t test, ****P<0.0001.

Since antigen is depleted from the surface after mechanical interaction, we aimed to validate that the decrease in early T cell activation relied on changes in the τ_F_ – not only a decrease in antigen density on the surface. By decreasing the concentration of FUSE probe incubated on the surface by half, we demonstrated that pYZAP70 levels were unaffected by a 60% reduction in antigen density (**Figure S7**). These results, along with our previous results (**Figure 3F**) showing that >40% of antigen remains underneath cells 15 min after cell engagement, indicate that potential fluctuations in antigen availability caused by FUSE are not responsible for changes in T cell activation shown in **Figure 4**. Thus, we conclude that perturbation in T cell signaling is dictated by the length of tension duration allowed by FUSE probes, which ultimately governs the number of mechanical engagements TCRs experience.

To further validate this conclusion, we investigated the impact of decreasing the probe density down to single molecule antigen density as well as increasing the trigger force threshold of FUSE probes. Since serial engagement is more pronounced at low antigen density, we hypothesized that decreasing τ_F_ would also have greater impact on T cell activation at low ligand density.^3^ Thus, we incubated 40,000 times less FUSE probe (5 pM instead of 200 nM) to achieve single molecule probe density, which resulted in a 400-fold depletion of antigen on the surface compared to our initial assay (**Figure S8**). Since T cells fail to spread on surfaces presenting low levels of antigen, we also anchored dimeric ICAM-1 along with pMHC-FUSE probes to promote adhesion (**Figure 4D** and **S9**). Using these low antigen density-ICAM surfaces, we examined pYZAP70 expression in cells limited to a τ_F_ of 1.9 min. Interestingly, we found that pYZAP70 levels decreased by 23% of that observed in the no enzyme control, which was a more pronounced difference than what was observed in the initial “high” antigen density experiments (**Figure 4E** and **4F**). This result agrees with previous serial engagement literature, as serial mechanical engagement is also more influential at low antigen densities.

Next, we created a FUSE probe with an F_1/2_ of 17.0 pN to test signaling in cells interacting with probes that are less likely to be unfolded and cleaved. We have previously shown that OT-1 cells can easily unfold hairpins presenting N4 pMHC with an F_1/2_ of 12 pN, but not 19 pN.^21^ Therefore, we were expecting background-level signal associated with the 17.0 pN FUSE probe unfolding. However, we were able to detect tension signal using the locking strategy, albeit much lower than tension signal reported with the 7.1 pN probe (**Figure S10**). When τ_F_ was limited to 1.9 min using high density of the 17.0 pN probe, we observed a 12% decrease in pYZAP70 with a lower statistical significance than the same experiment with the 7.1 pN FUSE probe (p < 0.05 for 17 pN, p < 0.0001 for 7.1 pN) (**Figure S11**).

## Discussion

Previous work has generated evidence supporting the TCR mechanosensor and serial engagement models as mechanisms of TCR triggering. Interestingly, the serial engagement model is primarily understood from the perspective of a series of connected reactions that are under kinetic control such that repeated binding of an antigen by different TCRs in proximity enhances and augments activation to allow for ultrahigh sensitivity.^38-40^ The mechanisms mediating the mechanosensor model are proposed to include kinetic mechanisms, such as catch bonds, as well as structural mechanisms where forces expose cryptic binding sites to the antigen.^11, 41-43^ These models are not necessarily mutually exclusive, and as such it is plausible that both models contribute to the sensitivity and specificity of the TCR. That said, there is little experimental evidence supporting the notion that the TCR takes advantage of both processes simultaneously, which we describe as serial mechanical engagement.

Our FUSE method provides a unique way to test the contributions of serial mechanical engagement, as our probes provide the experimenter with control over the length of time an antigen remains tethered to a surface once it is mechanically sampled with a precise pN tug. This design differs from probes that instantaneously rupture in response to force, such as DNA tension gauge tethers (TGTs), as these probes rupture and disrupt forces as soon as force is exerted onto an antigen.^44^ Experiments using the TGT binary response to force disruption provide evidence for the mechanosensor model but cannot explore the contributions of serial engagement on mechanosensing. Our results coupled with the results gathered using DNA TGTs show a distinct causal relationship between the magnitude of applied tension and the total duration of tension of the TCR-pMHC bond on T cell signaling (**Figure 5**). T cells plated on 12 pN OVA TGTs displayed a decrease of ∼60% in proximal kinase signaling compared to cells plated on 56 pN TGTs that cannot be opened by TCR forces.^21^ By definition, the TGT threshold is estimated at 2 s, and hence the 60% dampening in pY-ZAP70 levels for the 12 pN TGT represents the upper limit of how force duration reduces activation. This prior literature can be compared to our findings, wherein τ_F_’s of 1.9 min and 5.8 min led to a decrease in mean pYZAP70 of 15% and 8%, respectively, compared to an infinite τ_F_ probe. These findings demonstrate that the length of time that TCRs can mechanically interact with their antigens or, the number of successful mechanical interactions, dictates the level of activation achieved by the T cell. Importantly, our controls showing the change in ligand density on surface and the length of time that ligands persist under cells after cleavage further corroborate the conclusion that tuning the duration of serial mechanical engagement is responsible for the change in T cell signaling observed.

**Figure 5.**
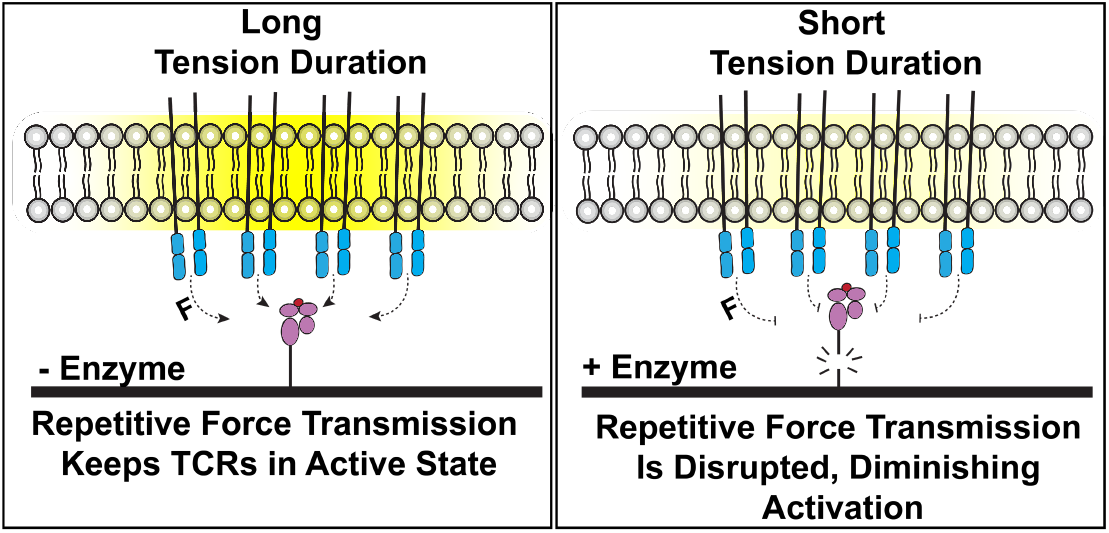
Schematic summarizing the correlation between activation of cells and the maximum TCR-pMHC tension duration afforded by FUSE probes.

Another key finding from this work is the exaggerated depletion of pYZAP70 that is observed when the τ_F_ is decreased on surfaces presenting low antigen density. This experiment most accurately depicts the physiological T cell-antigen presenting cell interface, as T cells encounter remarkably few stimulatory pMHCs on the surface of antigen presenting cells as they initiate their cytotoxic functions. Previous work has demonstrated that the reliance on serial engagement for TCR triggering is most pronounced when few agonist pMHCs are available to interact with a T cell.^3^ This observation is mirrored in our results, as the depletion of pYZAP70 detected in the lowest τ_F_ condition was reduced by 23% when the antigen density was decreased by 400-fold down to ∼1 antigen/μm^2^. Additionally, we found that limiting the τ_F_ of *F* >17.0 pN, which approaches the maximum magnitude TCR-pMHC force, resulted in a 12% decrease in pYZAP70 levels with τ_F_ reduced to 1.9 min (p < 0.05). The reduced impact on proximal kinase activity is most likely because interactions inducing a higher magnitude of force are less frequent, and thus fewer FUSE probes are mechanically triggered and cleaved. Therefore, serial mechanical engagement is likely operating across a range of forces but with lower frequency at *F* >17.0 pN.

The relationship between the τ_F_ of the TCR-pMHC interaction and T cell activation potential illustrated by FUSE is well supported by previous work using single-cell force spectroscopy.^16, 45^ While it has been well established that TCR-pMHC bond kinetics under force dictate the T cell’s functional response^11, 16, 17, 46, 47^, Pryshchep et al. demonstrate that the duration of cyclic, serially applied force onto TCRs also influence the activation potential of the cell.^45^ Here, they demonstrate that intracellular calcium levels are ∼20% lower in T cells that have encountered antigens serially applying forces for ∼2 minutes compared to calcium levels observed after 10 minutes of cyclic antigen engagement. Interestingly, little flux in calcium was observed when no forces were applied during a 10 min interaction between an antigen-decorated red blood cell and T cell. While FUSE probes limit the duration of serial TCR-mediated forces instead of ligand-mediated forces, limiting this duration reduced T cell activation by 15-23% depending on the ligand density, which agrees very well with the 20% reduction indicated in previous work. Our work further illustrates the necessity of serial mechanical engagements between TCRs and pMHCs, as the serial application of intrinsic (T cell-generated force) and extrinsic forces (applied by the experimenter) both lead to an increased T cell response over time. Moreover, the TCR-pMHC complex studied here has a bond lifetime ranging ∼0.1-1 s and thus our findings tuning force duration to 1.9 min demonstrates the importance of the integral of accumulated mechanical events rather than the outcome of single mechanical encounters in tuning T cell activation.^16^ This conclusion is in line with the notion of accumulated catch bonds triggering T cell activation as described by Zhu, Evavold and colleagues.^16, 45^

Ultimately, the development of FUSE probes has generated evidence for a link between two of the most prominent models used to explain T cell activation. While this design was able to investigate the connection between mechanosensing and serial engagement, optimization of FUSE could lead to further elucidation of the mechanisms of T cell activation. One limitation of the current design is that the lowest τ_F_ that we were able to achieve was still in the minute time regime, while TCR-pMHC interactions occur in the second to subsecond regime. To enhance this rate of disruption, future generations of FUSE probe could include a locking strand-restriction endonuclease conjugate that should ensure rapid enzymatic cleavage once the locking strand binds to the open hairpin. Another future direction for this project would be to use supported lipid bilayers to anchor FUSE probes instead of a glass substrate. This will allow antigens to freely diffuse on the surface, which mimics their physiological mobility on the surface of antigen presenting cells. Aside from T cell mechanobiology, FUSE could also be used to study the effects of tension duration on any mechanosensitive receptor of interest such as Notch, cadherins, and integrins. In fact, the highly tunable design of this probe provides a sturdy framework for multiple FUSE probes with different recognition sequences to be used to study the role that tension duration plays in a variety of receptors and co-receptors simultaneously.

## ASSOCIATED CONTENT

### Supporting Information

The Supporting Information is available free of charge on the ACS Publications website.

Materials and Methods, Supplementary Equations 1-4, Supplementary Figures 1-11(doc)

### AUTHOR INFORMATION

Present Addresses

Jhordan Rogers – *Department of Chemistry, Emory University, 1515 Dickey Drive, Atlanta, Georgia, 30322, USA*

Rong Ma – *Departments of Molecular and Cellular Physiology and Structural Biology, Stanford University School of Medicine, Stanford, CA 94305, USA*.

Yuesong Hu *– Department of Chemistry, Emory University, 1515 Dickey Drive, Atlanta, Georgia, 30322, USA*

## Author Contributions

All authors have given approval to the final version of the manuscript.

## Notes

The authors declare no conflicts of interest.

## ACKNOWLEDGMENT

The work was supported by NIH Grants RM1GM145394 and R01AI172452. Y.H. was supported by the National Cancer Institute Predoctoral to Postdoctoral Fellow Transition Award (F99CA274690). We would also like to thank the NIH Tetramer Facility for pMHC ligands.

## ABBREVIATIONS

TCR: T cell receptor
pMHC: peptide-major histocompatibility complex
FUSE: force-induced site-specific enzymatic cleavage
ZAP70: zeta-chain-associated protein kinase 70
N4: ovalbumin-derived SIINFEKL peptide antigen
ICAM-1: intercellular adhesion molecule-1
TGT: tension gauge tether.

## Notes

### Competing Interest Statement

The authors have declared no competing interest.

