## Supporting Information for "Force-induced site-specific enzymatic cleavage probes reveal that serial mechanical engagement boosts T cell activation"

**Affiliations:**
[1]Department of Chemistry, Emory University, 1515 Dickey Drive, Atlanta, Georgia, 30322, USA
[2]Wallace H. Coulter Department of Biomedical Engineering, Georgia Institute of Technology and Emory University, Atlanta, Georgia, 30332, USA

### Supplementary Note

FUSE DNA probes utilize the well-characterized mechanical unfolding of the DNA hairpin.[1, 5, 6] At thermodynamic equilibrium, the force required to unfold a DNA hairpin is defined as the $F_{1/2}$, which can be calculated using **Equation S1**:

$$F_{1/2} = \frac{\Delta G_{unfold} + \Delta G_{stretch}}{\Delta x}, \qquad \text{( S1 )}$$

where $\Delta G_{unfold}$ is the free energy of unfolding without force and $\Delta G_{stretch}$ is the free energy of stretching the ssDNA from its folded origin. This can be calculated using the worm like chain model in **Equation S2**:

$$\Delta G_{stretch} = \left(\frac{k_B T}{L_p}\right)\left(\frac{L_0}{4\left(\frac{1-x}{L_0}\right)}\right)\left[3\left(\frac{x}{L_0}\right)^2 - 2\left(\frac{x}{L_0}\right)^3\right], \qquad \text{( S2 )}$$

where $L_p$ is the persistence length of ssDNA (~1.3 nm), $L_0$ is the contour length of ssDNA (0.63 nm / nucleotide), x is the hairpin extension from equilibrium by using 0.44 x [(# of nucleotides in hairpin)-1]. Δx in **Equation S1** can be found by subtracting 2 nm from x, as this distance corresponds to the width of the DNA hairpin stem duplex where the unfolding process begins.

Assuming pseudo first-order kinetics, rate of hybridization between the locking strand and hairpin was determined by fitting the percentage of maximum locking strand binding versus time with the one phase association function in **Equation S3**:

$$Y = Y_0 + (Y_\infty - Y_0)(1 - e^{-kx}), \qquad \text{( S3 )}$$

where $Y_0$ is the percentage of maximum locking strand binding at x=0, $Y_\infty$ is the percentage of maximum binding as x approaches infinity, and k is the rate constant expressed as $min^{-1}$.

The rate of enzymatic cleavage of locked FUSE DNA probes was determined by fitting the one phase exponential decay function in **Equation S4** to normalized fluorescence intensity versus time plots for each concentration of AseI.

$$Y = (Y_0 - Y_\infty)e^{-kx} + Y_\infty \qquad \text{( S4 )}$$

Here, $Y_0$ is the normalized fluorescence intensity at x=0, $Y_\infty$ is the fluorescence intensity as x approaches infinity, and k is the rate constant expressed as $min^{-1}$. The average lifetime of locked FUSE DNA probes ($\tau_F$) is the reciprocal of k and the half-life ($t_{1/2}$) is calculated by dividing ln(2) by k.

### Materials and Methods

#### Oligonucleotides

All oligonucleotides were custom synthesized by Integrated DNA Technologies (Coralville, IA). **Table S1** includes the names and sequences for all oligonucleotides used in this work.

.

**Table S1. List of oligonucleotides.**

| Strand ID | Description of strand | Sequence (5' to 3') |
| --- | --- | --- |
| 7.1 pN FUSE DNA hairpin (Long Stem) | Hairpin containing AseI recognition site with $F_{1/2}$ of 7.1 pN | GTG AAA TAC CGC ACA GAT GCG TTT GTA TAA ATA AAT GAT TAA TGC ATT TAT TTA TAC TTT AAG AGC GCC ACG TAG CCC AGC |
| 4.7 pN FUSE DNA hairpin (Short Stem) | Hairpin containing AseI recognition site with $F_{1/2}$ of 4.7 pN | GTG AAA TAC CGC ACA GAT GCG TTT GTA TAA ATG ATT AAT ACA TTT ATA CTT TAA GAG CGC CAC GTA GCC CAG C |
| Traditional 4.7 pN Hairpin | Hairpin without AseI recognition site with $F_{1/2}$ of 4.7 pN | GTG AAA TAC CGC ACA GAT GCG TTT GTA TAA ATG TTT TTT TCA TTT ATA CTTTAA GAG CGC CAC GTA GCC CAG C |
| 17 pN FUSE DNA hairpin | Hairpin containing AseI recognition site with $F_{1/2}$ of 17 pN | GTG AAA TAC CGC ACA GAT GCG TTT CGG GCC GGC GCG CGG ATT AAT CCG CGC GCC GGC CCG TTT AAG AGC GCC ACG TAG CCC AGC |
| A21B | Ligand presenting strand of hairpin probe | /5AmMC6/ - CGC ATC TGT GCG GTA TTT CAC TTT - /3Bio/ |
| Cy3B strand | Cy3B labeled A21B strand | Cy3B - CGC ATC TGT GCG GTA TTT CAC TTT - /3Bio/ |
| BHQ-2 strand | Anchor and quencher strand of hairpin probe | /5-ThioC6-5/ - TTT GCT GGG CTA CGT GGC GCT CTT - /3BHQ_2/ |
| 15mer – short stem FHP | 15 nt locking strand for 4.7 pN FUSE DNA hairpin | ATT AAT CAT TTA TAC |
| 17mer – short stem FHP | 17 nt locking strand for 4.7 pN FUSE DNA hairpin | GTA TTA ATC ATT TAT AC |
| 17mer – THP | 17 nt locking strand for traditional 4.7 pN hairpin | GAA AAA AAC ATT TAT AC |
| 21mer – long stem FHP | 21 nt locking strand for 7.1 pN FUSE DNA hairpin | GCA TTA ATC ATT TAT TTA TAC - /3AmMO/ |
| Atto647N 21mer – long stem FHP | 21 nt locking strand for 7.1 pN FUSE DNA hairpin labeled with Atto647N | GCA TTA ATC ATT TAT TTA TAC - Atto647N |

| | | |
|---|---|---|
| 22mer – 17 pN FHP | 22 nt locking strand for 17.0 pN FUSE DNA hairpin | GAT TAA TCC GCG CGC CGG CCC G |
| Unstructured 7.1 pN FHP | Docking site for 21mer locking strand to mimic 7.1 pN FHP opening | GTA TAA ATA AAT GAT TAA TGC CCA GCG TGA T - /3ThioMC3-D/ |

**Reagents**
(3-Aminopropyl)triethoxysilane was purchased from Acros (Cat# AC430941000, Pittsburgh, PA). LA-PEG-SC (Cat# HE039023-3.4K) and mPEG-SC (Cat# MF001023-2K) were purchased from Biochempeg (Watertown, MA). Sulfo-NHS acetate (Cat# 26777) and phospho-ZAP70/Syk polyclonal antibody (Cat# PA5-17815) were purchased from Thermo Fisher Scientific (Waltham, MA). Goat anti-rabbit IgG Alexa Fluor 488 (Cat# ab150077) was purchased from Abcam (Cambridge, UK). Custom synthesized 8.8 nm diameter tannic acid modified gold nanoparticles were obtained from Nanocomposix (San Diego, CA). Ethanol (Cat# 459836), hydrogen peroxide (Cat# H1009), bovine serum albumin (BSA) (Cat# 10735078001), Atto647N NHS ester (Cat# 18373-1MG-F), 10 kDa MWCO amicon ultra-0.5 centrifugal filter units (Cat# UFC501096), and Hank's balanced salts (H8264) were purchased from Sigma Aldrich (St. Louis, MO). Dulbecco's phosphate-buffered saline (DPBS, Cat# 21-031-CM), and Dulbecco's Modified Eagle's Medium (DMEM, 10-013-CV) were acquired from Corning. Cy3B NHS ester (Cat# PA63101) was purchased from GE Healthcare (Pittsburgh, PA). NTA-SAM reagent was purchased from Dojindo Molecular Technologies (Rockville, MD). Red blood cell lysis buffer (Cat# 00-4333-57), and biotinylated anti-mouse CD3ε (Cat# 13-0031-82). Biotinylated pMHC ovalbumin (SIINFEKL) was obtained from the NIH Tetramer Core Facility at Emory University, which has been described in detail in our previous work (4). Teflon racks (Cat# C14784) were obtained from Thermo Fisher Scientific. P2 size exclusion gel (Cat#1504118) was purchased from Bio-Rad (Hercules, CA). 3 mL syringes were purchased from BD bioscience (San Jose, CA). Cell strainers (Cat# 15-1100) were bought from Biologix (Shandong, China). Nanosep MF centrifugal devices (Cat# ODM02C35) with bio-inert membrane were bought from Pall laboratory (Port Washington, NY). Midi MACS (LS) startup kit (Cat# 130-042-301) (separator, columns, stand), and mouse CD8+ T cell isolation kit (Cat# 130-104-075) were purchased from Miltenyi Biotec (Bergisch Gladbach, Germany). AseI restriction endonuclease with NEBuffer 3.1 (Cat# R0526S) was purchased from NEB (Ipswich, MA).

**Cells**
The OT-1 CD8+ T cell is a well-established system to study T cell biology. OT-1 transgenic mice were housed and bred in the Division of Animal Resources Facility at Emory University under the Institutional Animal Care and Use Committee protocol PROTO201800239. Naïve OT-1 T cells that express the CD8 co-receptor and specifically recognize chicken ovalbumin epitope 257–264 (SIINFEKL) were isolated and enriched from the spleen of a sacrificed mouse using MACS separation with the CD8+ T cell isolation kit according to the manufacturer's instructions. DPBS buffer supplemented with 0.5% BSA and 2 mM EDTA was used as a buffer for the purification process. The purified CD8+ naïve OT-1 cells were kept in Hank's balanced salt solution (H8264) at 4 °C before resuspending in NEBuffer 3.1 directly before imaging.

**Equipment**
The major equipment that was used in this study includes: Barnstead Nanopure water purifying system (Thermo Fisher), High-performance liquid chromatography (Agilent 1100), Nanodrop 2000 UV-Vis Spectrophotometer (Thermo Scientific), Inverted microscope system (Nikon Eclipse Ti) equipped with an EMCCD, total internal reflection fluorescence (TIRF) microscopy components TIRF 488 nm, TIRF 561 nm, TIRF 647 nm, a perfect focus system for maintaining focus during timelapse imaging, and reflection interference contrast microscopy (RICM).

**Oligonucleotide-dye conjugation**
Oligonucleotide-dye conjugates (Cy3B strand, 21mer locking strand) were prepared by coupling the terminal amine on the DNA strand with activated NHS-ester of the organic dye. Briefly, 50 μg (an excess quantity) of NHS dye was dissolved in 10 μL of DMSO and reacted with 10 nmol of oligonucleotide in 1×PBS with 0.1 M $NaHCO_3$ for 1 h at room temperature. After the reaction, byproducts, salts, and unreacted dye in the mixture were removed by P2 gel filtration with Nanosep MF centrifugal devices at 14000 rpm for 1 min. The product was further purified

by reverse-phase HPLC with an Agilent AdvanceBio Oligonucleotide C18 column (653950-702, 4.6 x 150 mm, 2.7 μm). The mobile phase A: 0.1 M TEAA and B: ACN were used for a linear gradient elution of 10-100% B over 50 min at a flow rate of 0.5 mL/min. The concentration of the strands was determined by UV-Vis using a Nanodrop instrument.

**His-tagged dimeric GFP-ICAM-1 expression**

GFP-ICAM-1 constructs were expressed and purified using the same protocol described in (7). HEK293FT cells for lentivirus production were maintained in complete DMEM supplemented with 10% FBS, penicillin G (100 U/mL), and streptomycin (100 μg/mL) at 37 °C with 5% CO2. Lentivirus particles were produced with HEK293FT cells by cotransfection of the pLEX transfer plasmid encoding the soluble dimeric ICAM-1 sequence with the 2nd generation packaging plasmids pMD2.G and psPAX2 (gifts from Didier Trono, Addgene plasmid #12259 and #12260) using linear polyethylenimine (MW = 25,000). The particles were harvested from the supernatant 48-72 h post-transfection, filtered and concentrated into ~200 μL in DMEM with 20% FBS by ultracentrifugation and stored at - 80 °C before use. For lentiviral transduction, ~20,000 TB-15 cells (a variant of HEK293T cells stably expressing BirA biotinylating enzyme, a gift from Prof. John Altman and Dr. Richard Willis from the NIH Tetramer Core Facility at Emory University) were seeded onto a 96 well cell culture plate. On the next day, ~50 μL of concentrated lentivirus particles were added to the cells. After 6 h of infection, the media was exchanged to complete DMEM. Transduced cells were expanded to appropriate density before adaptation to suspension culture in a shaking incubator where the cells were maintained in FreeStyle 293 expression media (8% CO2, shaking speed = 125 rpm). Soluble ICAM-1 was collected from the supernatant every 3 days and purified using Ni-NTA agarose affinity resin (Thermo Fisher Scientific). ICAM-1 quality was confirmed using SDS-PAGE and fluorescence imaging. The concentration of the recombinant ICAM-1 protein was quantified using Nanodrop and adjusted to 1 mg/mL and stored at -80 °C before use.[7]

**Preparation of hairpin probes on gold nanoparticle surfaces**.

Gold nanoparticle hairpin probe surfaces were prepared as described previously (4) and as illustrated in Fig. S1. Note that we have made slight modifications to previously published methods. Briefly, glass slides were placed in a Teflon rack and rinsed with 40 mL of Nanopure water 3 times. Then the rack holding the slides was placed in a beaker with 1:1 (v/v) solution of ethanol:Nanopure water and sonicated for 20 min. After sonication, the slides were washed 6 times with 40 mL of Nanopure water to remove any remaining organic solvents. 9 Next, 40 mL of freshly made piranha solution (3:1 mixture (by volume) of $H_2SO_4$ and $H_2O_2$, CAUTION: highly explosive if mixed with organics) was prepared. The rack carrying slides was submerged in piranha solution for 30 minutes. After 30 min, the slides were washed extensively with 40 mL of Nanopure water 6 times, followed by another 3 washes with 40 mL of ethanol, until water was removed. Next, slides were incubated in 3% APTES in 40 mL of ethanol for 1 h at room temperature, after which the surfaces were washed with 40 mL of ethanol 3 times and baked in oven at 80 °C for 20 min. Then, the amine-modified glass coverslips were placed in petri dishes lined with parafilm with 300 μL of 0.5% w/v lipoic acid-PEG-NHS and 2.5% w/v mPEG-NHS in 0.1 M NaHCO3 on each slide for 1 h at room temperature. Subsequently, the samples were rinsed with Nanopure water and incubated with 0.1 M $NaHCO_3$ containing 1 mg/mL of sulfo-NHS acetate for 30 min in order to passivate unreacted amine groups. Gold nanoparticles (8.8 nm, tannic acid modified) at 20 nM were subsequently added to surfaces for 30 minutes after extensive washes with Nanopure water. Meanwhile, the hairpin, Cy3B, and BHQ2 strands that form the molecular tension probe constructs were mixed and annealed at a ratio of 1.1:1:1 in 1 M NaCl at concentrations ranging from 5 pM-300 nM. After annealing, an additional 2.7 μM of BHQ2 strand was added to the DNA solution for additional quenching. After 30 min of surface incubation with gold nanoparticles at room temperature and three washes with Nanopure water, the annealed DNA solution containing tension probes and quencher was added to surfaces and allowed to be immobilized overnight at 4°C. After washing excess probes away with PBS on the second day, 40 μg/mL of streptavidin in PBS was added to the surfaces and incubated for 30 min. After washing slides with PBS, 40 μg/mL biotinylated antiCD3ε or N4 pMHC was allowed to bind to tension probes via biotin-streptavidin interactions for 30 min at room temperature in PBS. Finally, the slides were rinsed with PBS, assembled in imaging chambers, and immediately used for imaging.

**Co-presenting ligands on surfaces modified with MTFM probes**

Gold nanoparticles at 10 nM were added to surfaces modified with mPEG and lipoic acid PEG and allowed to react for 30 min. After washing 3 times with Nanopure water and twice with ethanol, 100 μL 2 mM NTA-SAM in ethanol was added to each surface and incubated at room temperature for 1 h in a sealed petri dish. The excess NTA-SAM was washed away with ethanol 3 times, followed by another 3 washes with Nanopure water. Gold nanoparticles at 20 nM were then added to surfaces again and incubated at room temperature for 30 min. Meanwhile the DNA tension probes were annealed as described previously. After hybridization, the probes were allowed to anchor on gold particles overnight at 4 °C. On the following day, 10 mM NiCl2 in PBS was added to surfaces for 10 min and washed away with PBS. Dimeric ICAM-1 with His-tag was immobilized on surfaces by incubating surfaces with 10 μg/mL solution in PBS. After washing with PBS, streptavidin and biotinylated pMHC N4 were added to surfaces sequentially at 40 μg/mL as previously described.

**Fluorescence imaging**
Epi fluorescence microscopy and total internal reelection fluorescence (TIRF) microscopy experiments were performed on a Nikon Eclipse Ti inverted microscope driven by the NIS Elements software. The microscope features an Evolve electron-multiplying charge-coupled device (Photometrics), an Intensilight epifluorescence source (Nikon), CFI Apo 100 X (numerical aperture 1.49) objective and 40 X (numerical aperture 0.75) objective (Nikon), and a total internal reflection fluorescence launcher with three laser lines: 488 (10 mW), 561 (50 mW), and 638 nm (20 mW). Methods use the 100x objective unless otherwise stated. This microscope also includes the Nikon Perfect Focus System, an interferometry-based focus lock which allows the capture of multi-point and time-lapse images without loss of focus. Imaging was performed using either Hank's balanced salt solution or NEBuffer 3.1. All imaging data was acquired at 37°C.

**Imaging in cell-free experiments**
Surfaces presenting hairpin tension probes were fabricated the same way as described previously but without streptavidin and ligands. Surfaces presenting the locked hairpin probe were prepared by adding 10x locking strand compared to hairpin probe concentration. Surfaces presenting unstructured sequence were prepared similarly, except for using the same amount of unstructured sequence instead of DNA tension probe construct. The surfaces were imaged and incubated with locking oligonucleotides ranging from 5–2000 nM and AseI concentrations ranging from 0.5 – 10.7 nM. Timelapse data was acquired at 3 xy-coordinates on each surface and averaged for processing after subtracting CCD background. All kinetics experiments were performed in NEBuffer 3.1 without cells and repeated 3 times.

**Mechanically selective hybridization and specific enzymatic cleavage**
Naïve CD8+ OT-1 cells were purified from a transgenic OT-1 mouse 1 h before imaging. Purified OT-1 cells and 200 nM of Atto647N labeled locking strand in NEBuffer 3.1 were added to surfaces presenting N4 pMHC. After 15 min, the cells were imaged, and wells were gently rinsed with 1xPBS to remove excess locking strand. 10.7 nM of AseI was then added in fresh NEBuffer 3.1 to track the loss of tension and locked signal associated with enzymatic cleavage (**Figure 3**). A timelapse video tracked the change in both signals every minute for 20 minutes. The mean fluorescence intensity under cells was subtracted by the mean fluorescence intensity of areas of the surface without cells (background signal) to determine the tension and locked signals. A traditional 4.7 pN hairpin without the recognition sequence for AseI was used as control to demonstrate the specificity of enzymatic cleavage.

**FUSE DNA assay and immunostaining**
$1x10^5$ naïve CD8+ OT-1 cells, 5 μM locking strand, and 0 – 10.7 nM AseI in 1xNEBuffer 3.1 were added to surfaces presenting N4 pMHC. After a 15 min incubation at 37°C, cells were gently washed with PBS and fixed with 4% formaldehyde in PBS for 10 min. The surfaces were gently washed with PBS, permeabilized in 0.1% Triton X-100 for 10 min, and then washed again with PBS. Subsequently, 2% BSA was added to the surfaces and incubated overnight at 4°C. On the next day, the surfaces were washed with PBS. Phospho-ZAP70/Syk polyclonal antibody (Cat# PA5-17815) was diluted to 1:100 in staining buffer (0.5% BSA and 0.01% Triton X-100 in PBS) and added to the surface for 1 h incubation at room temperature. After washing thoroughly with PBS, goat anti-rabbit IgG Alexa Fluor 488 (Cat# ab150077) secondary antibody was added at a dilution of 1:500. Surfaces were then washed with PBS before epifluorescence imaging with a 40x objective.

**Statistical analysis**

All experiments were conducted as at least three technical or biological replicates. Statistical significance was determined on GraphPad Prism using two-tailed student's *t* tests. Data are presented as bars with mean±SD or scatter plots with line representing mean

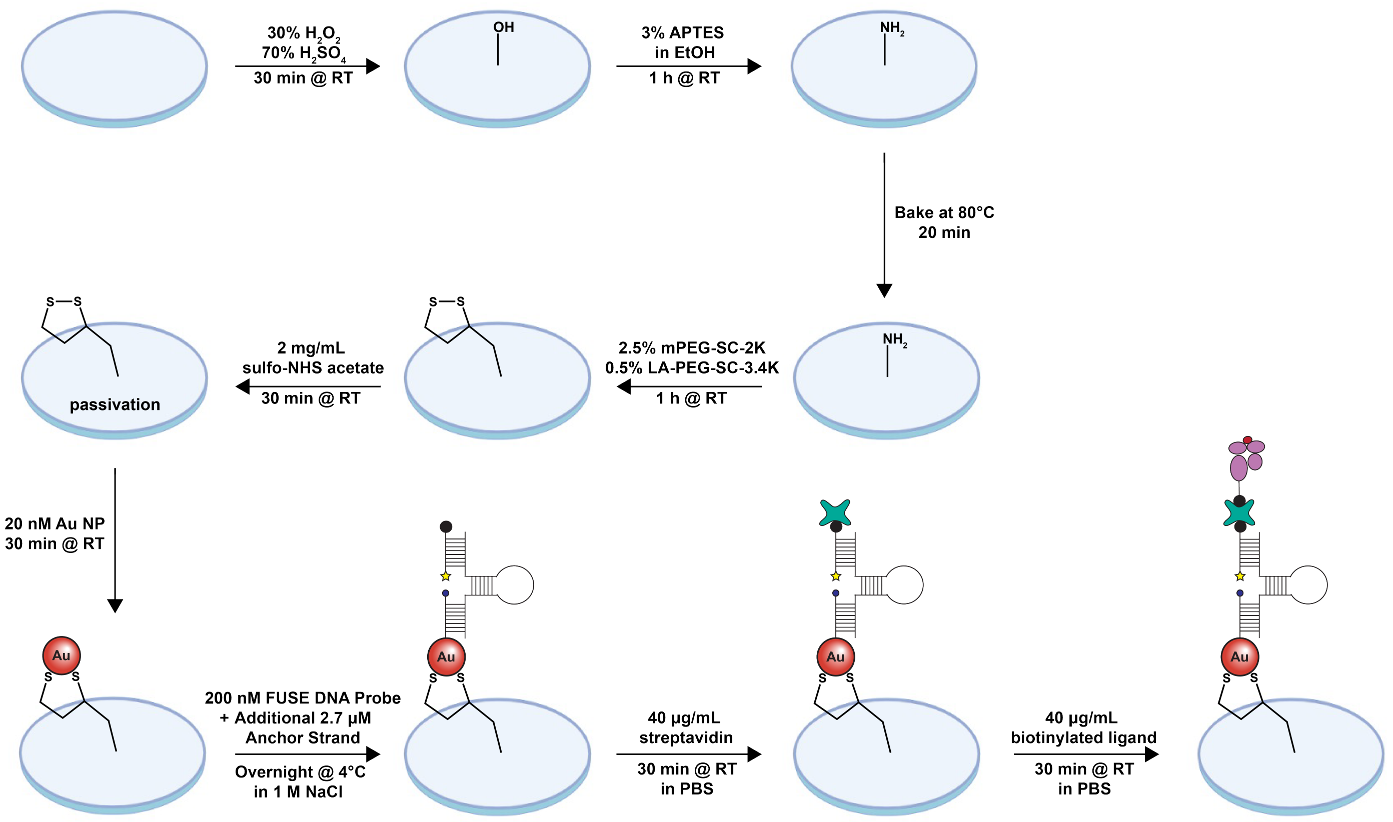

**Figure S1. DNA hairpin-gold nanoparticle surface preparation protocol.**

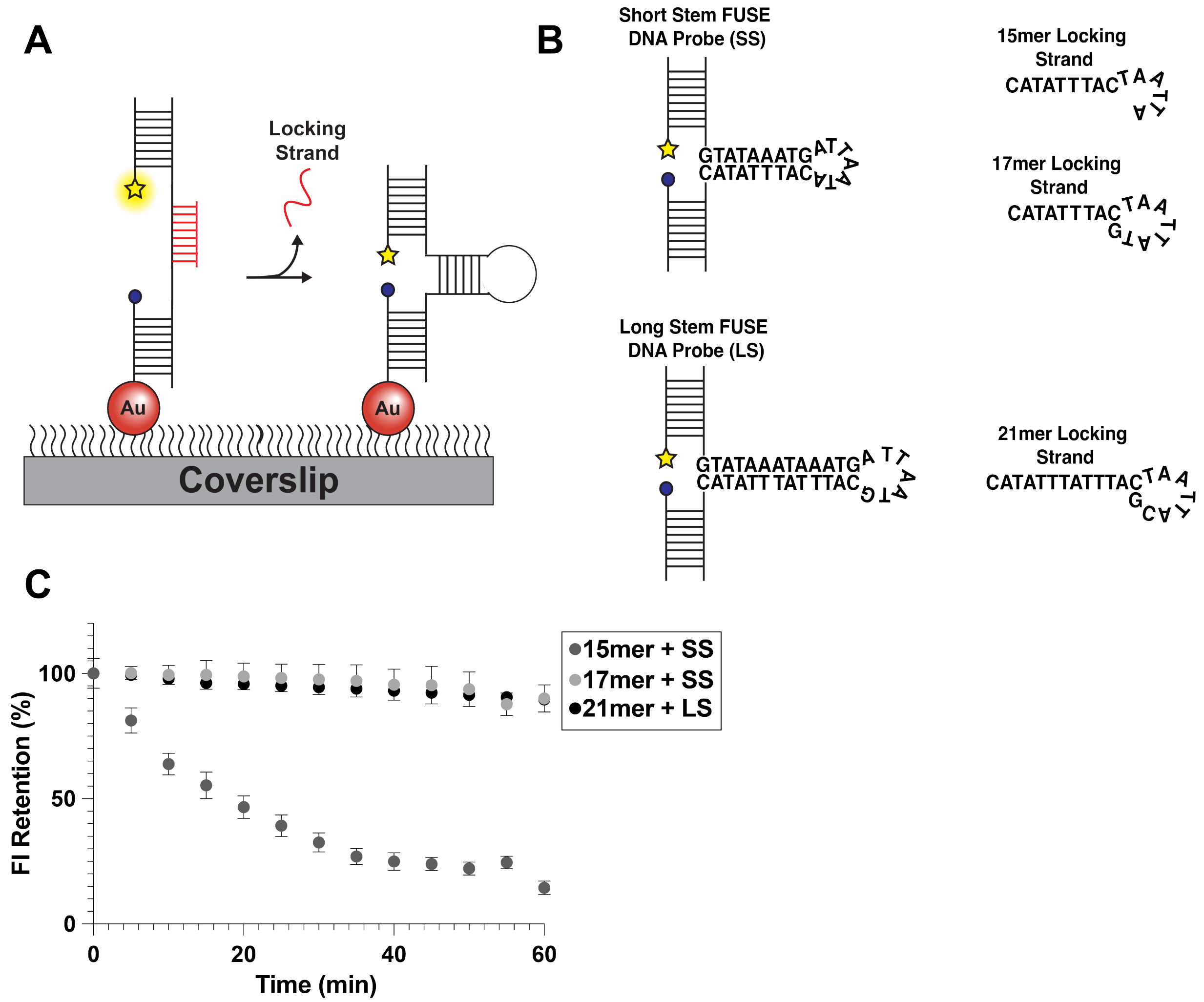

**Figure S2. FUSE DNA sequence screen.** A) Schematic showing the dissociation of the locking strand from the DNA hairpin as a result of low thermostability of the hairpin-locking strand duplex. B) FUSE DNA hairpin and locking strand sequences used to test thermostability of the DNA hairpin-locking strand complex. The short stem FUSE DNA probe (SS) is composed of a stem region 9-nt long with a 7-nt loop. The 15mer locking strand binds to the full top portion of the stem, as well as 6-nt in the loop ($T_m$ = 36.5°C, 0.1 mM NaCl and 10 mM $MgCl_2$, 1 μM DNA). The 17mer binds to the full top portion of the stem, the full loop, and one nucleotide on the bottom portion of the stem ($T_m$ = 42.5°C, 0.1 mM NaCl and 10 mM MgCl2, 1 μM DNA). The long stem FUSE DNA probe (LS) is composed of a stem region 13-nt long with a 7-nt loop, wherein the 3' adenine is replaced with a guanine. The 21mer binds to the full top portion of the, the full loop, and one nucleotide on the bottom portion of the stem ($T_m$ = 51.5°C, 0.1 mM NaCl and 10 mM $MgCl_2$, 1 μM DNA). C) Plot of the change in fluorescence intensity associated with hairpin closing after locking strand dissociation at 37°C versus time. The 15mer + SS displayed poor thermostability ($t_{1/2}$ = 19.6 min), while neither the 17mer + SS nor the 21mer + LS surfaces decreased by more than 10% over the course of the hour-long incubation at 37°C. We elected to use the 21mer + LS combination due to the higher $T_m$ and inclusion of the guanine in the loop, as we hypothesized that the incorporation of an additional guanine-cytosine base pair could improve the stability of the initial lock-hairpin duplex formation.[2, 3]

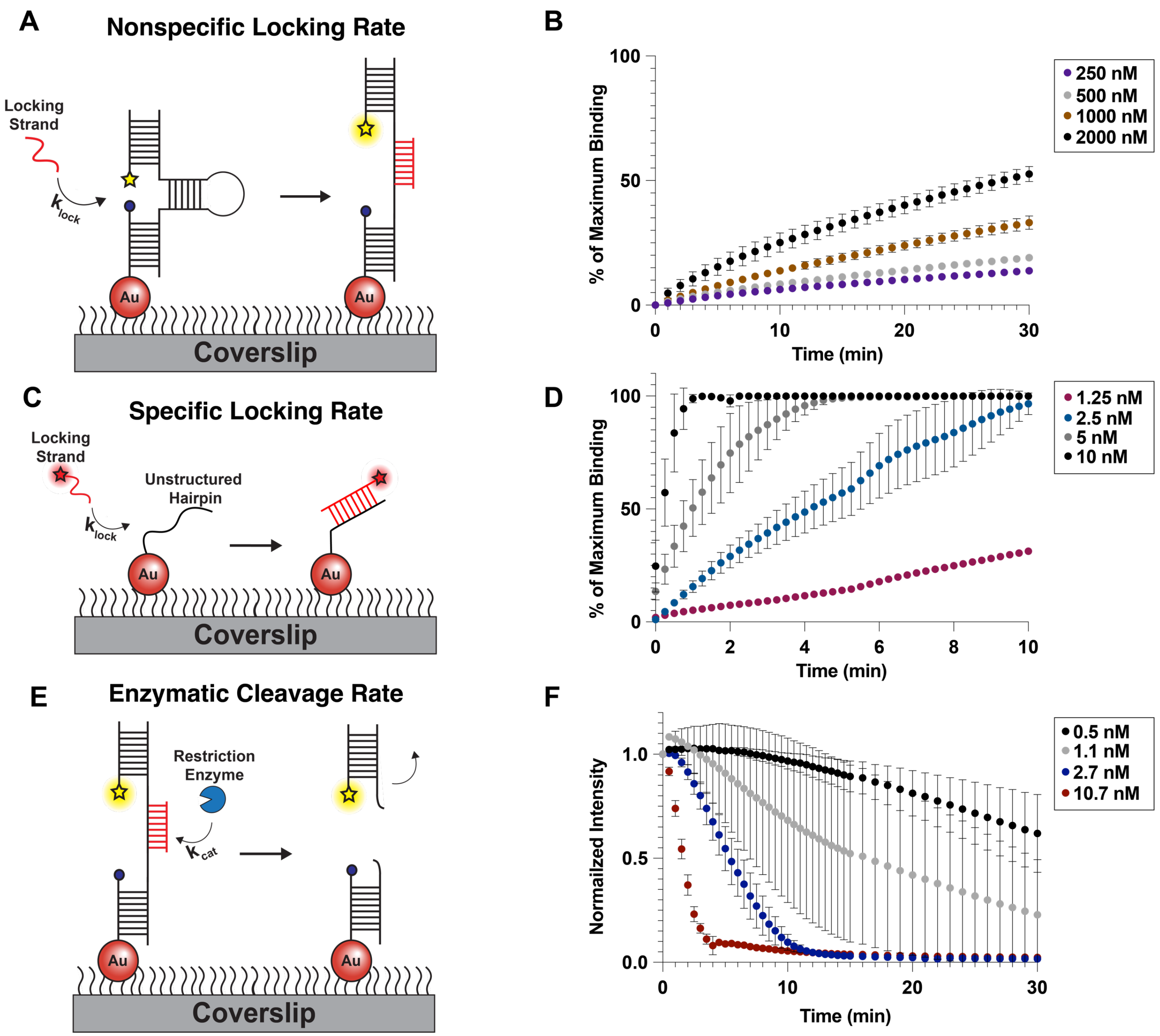

**Figure S3. Lock hybridization and enzymatic cleavage kinetics.** A) Schematic of experiment to quantify rate of nonspecific hybridization of locking strand (see also **Figure 2A**). B) The percentage of maximum locking strand binding to the closed hairpin versus time for locking strand concentrations ranging from 250 nM–2000 nM. C) Schematic of experimental design to quantify the rate of specific hybridization of locking strand to its exposed docking site (see also **Figure 2D**). D) Plot of the percentage of maximum locking strand binding to the unstructured hairpin versus time for locking strand concentrations ranging from 1.25 nM–10 nM. E) Schematic displaying the experimental setup to quantify the rate of enzymatic cleavage of locked hairpin (see also **Figure 2G**). F) Plot of the normalized surface intensity versus time to quantify the apparent rate of lock-hairpin duplex cleavage with AseI concentrations ranging from 0.5 nM–10.7 nM.

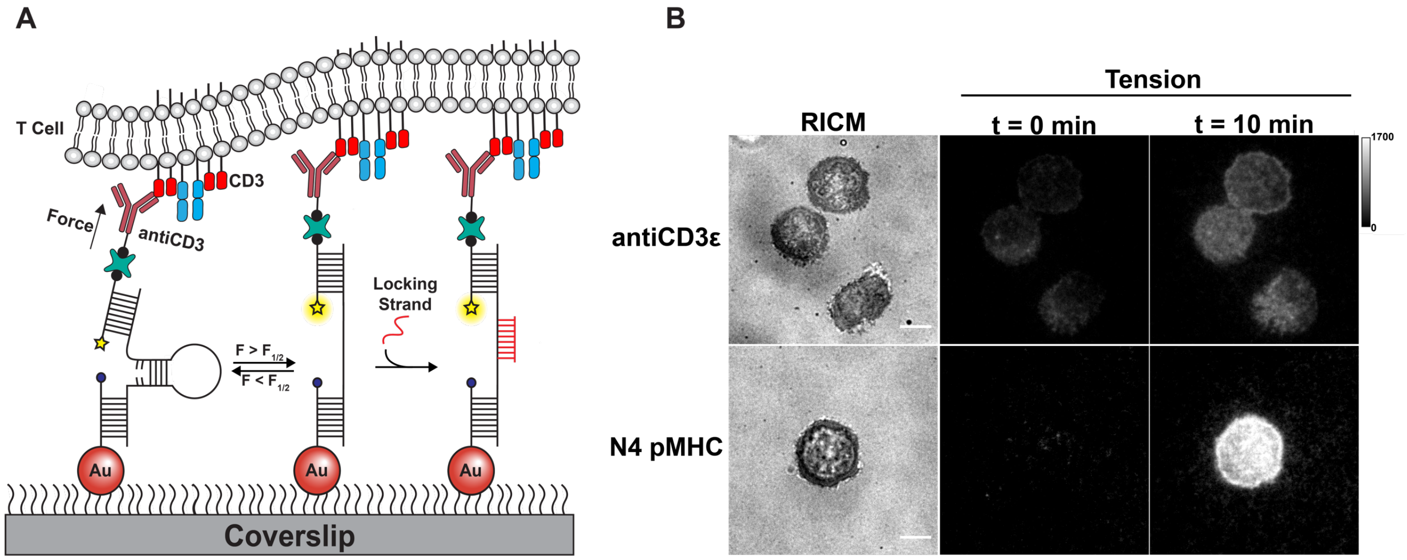

**Figure S4. Validation of opening and locking of 7.1 pN FUSE DNA probe.** A) Schematic depicting locking of FUSE DNA probes presenting antiCD3ε as force is exerted through CD3ε subunit of the TCR/CD3 complex. B) Representative images of cell spread area (RICM) and tension signal before (t = 0 min) and after (t = 10 min) adding 200 nM of the 21mer locking strand on surfaces presenting either antiCD3ε or N4 pMHC on 7.1 pN FUSE DNA probes. Tension signal is typically higher before adding locking strand on antiCD3ε surfaces compared to N4 pMHC surfaces due to the long-lived interactions between the antibody and antigen. Tension signal after incubating locking strand for 10 min is typically higher on surfaces presenting N4 pMHC compared to antiCD3ε due to the high mechanical sampling rate between the TCR and pMHC.[4]

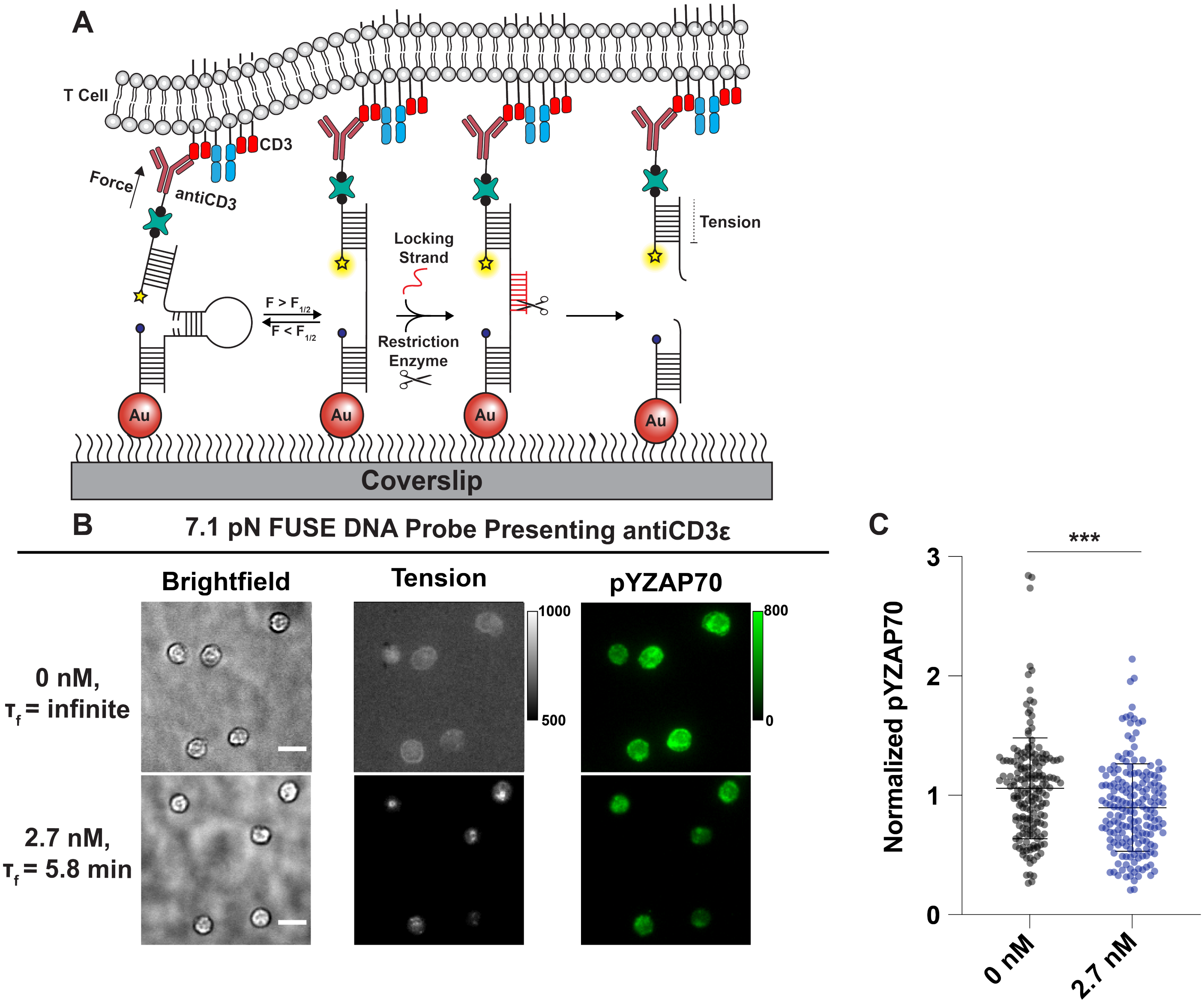

**Figure S5. Limiting $\tau_f$ of antiCD3ε-CD3ε interaction with 7.1 pN FUSE DNA probes.** A) Schematic of the enzymatic cleavage of FUSE DNA probes presenting antiCD3ε. FUSE DNA assay most likely prematurely terminates single antibody-antigen mechanical interactions rather than serial mechanical engagement in this case due to the long-lived interactions between antiCD3ε and CD3ε. B) Representative images of cells interacting with surfaces (brightfield), tension signal, and pYZAP70 signal after 15 min incubation with or without 2.7 nM AseI on surfaces presenting antiCD3ε on FUSE DNA probes. C) Comparison between the normalized pYZAP70 signal between the enzyme (mean norm. pY = 0.89) and no enzyme conditions (mean norm. pY = 1.05), n>180 cells. Statistical analysis was performed using Student's t test, ***P<0.001.

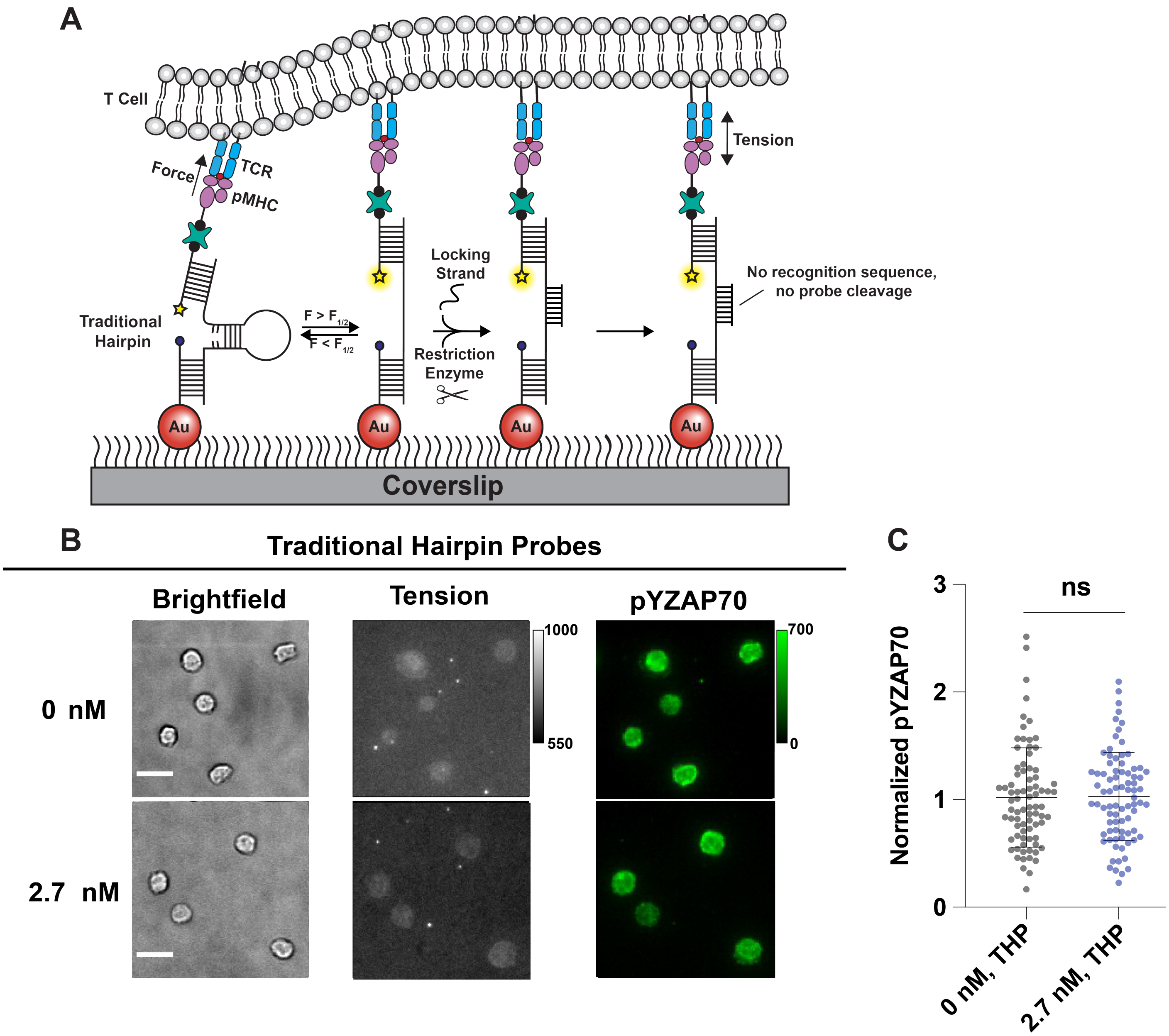

**Figure S6. Control to test role of specific enzymatic cleavage on pYZAP70 expression.** A) Schematic of the addition of AseI to probes that do not contain its recognition sequence. B) Representative images of cells interacting with surfaces (brightfield), tension signal, and pYZAP70 signal after 15 min incubation with or without 2.7 nM AseI on surfaces presenting N4 pMHC on traditional 4.7 pN hairpin probes. C) Comparison between the normalized pYZAP70 signal between the enzyme (mean norm. pY = 1.03) and no enzyme conditions (mean norm. pY = 1.02), n>80 cells. Statistical analysis was performed using Student's t test, ns = P>0.05.

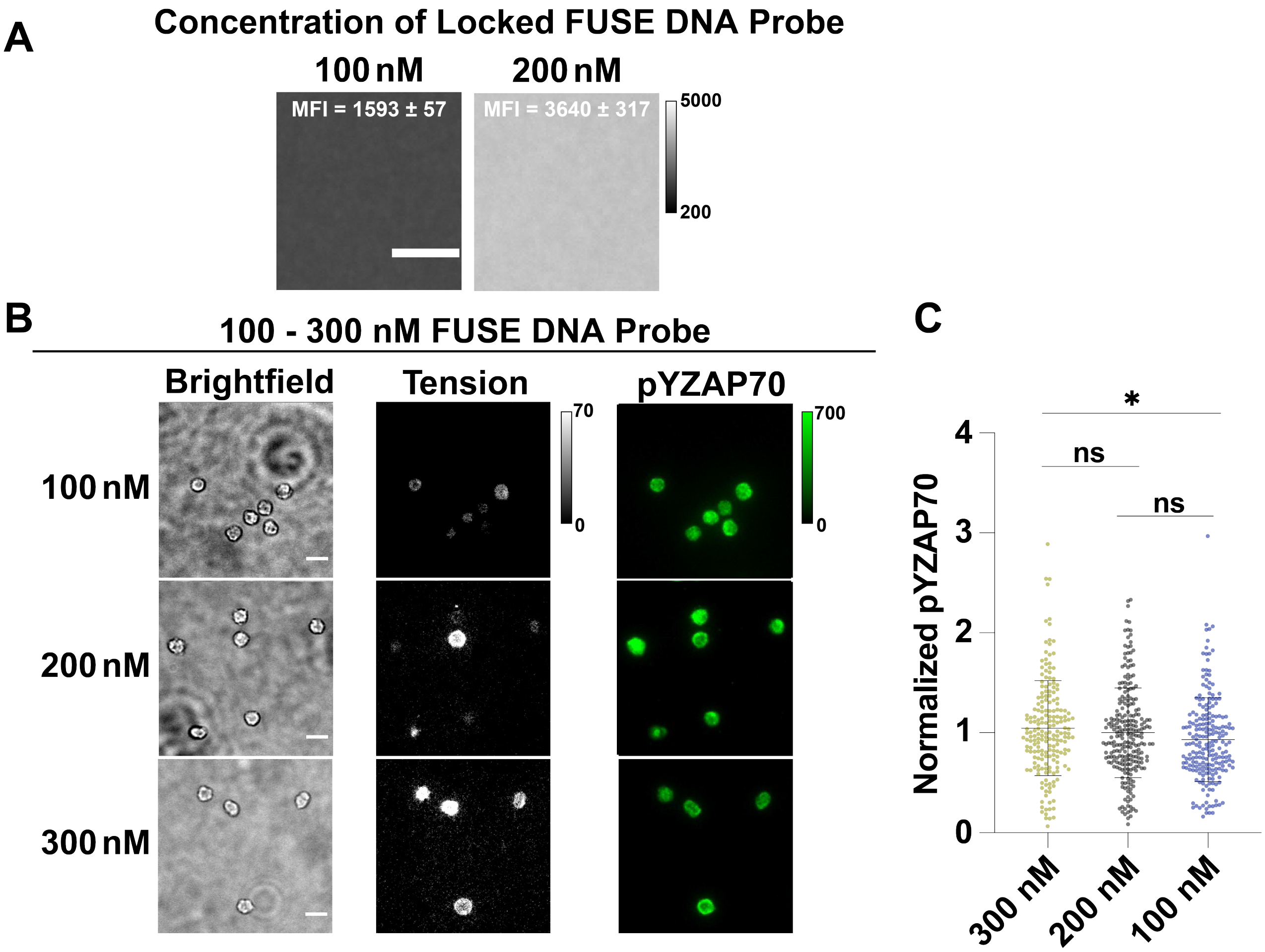

**Figure S7. Testing the effect probe density has on pYZAP70 expression.** A) Representative images of surfaces after incubating either 100 nM or 200 nM locked FUSE DNA probes. Using fluorescence intensity of these surfaces, the 100 nM incubation results in ~60% depletion in probe density on surface compared to the 200 nM incubation. B) Representative images of cells interacting with surfaces (brightfield), tension signal, and pYZAP70 signal after 15 min incubation on surfaces presenting N4 pMHC with a range of FUSE DNA probes densities. C) Comparison of the normalized pYZAP70 signal in cells interacting with surfaces incubated with 100 nM probe (mean norm. pY = 0.93), 200 nM probe (mean norm. pY = 1.00), and 300 nM probe (mean norm. pY = 1.04), n>80 cells. Statistical analysis was performed using Student's t test, ns = $P>0.05$ and $^{*}P<0.05$.

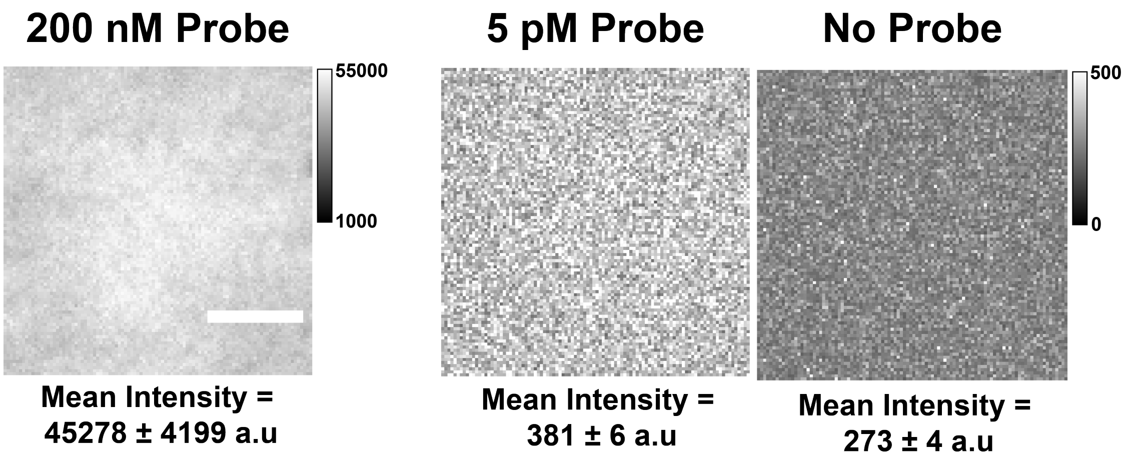

**Fold Decrease:**
**(381-273)/45278 = 0.0024 ± 0.0004**

**Figure S8. Quantification of relative decrease in probe density for single molecule assay.** Comparison between fluorescence intensity of surfaces after incubating 200 nM, 5 pM, or no locked FUSE DNA probe.

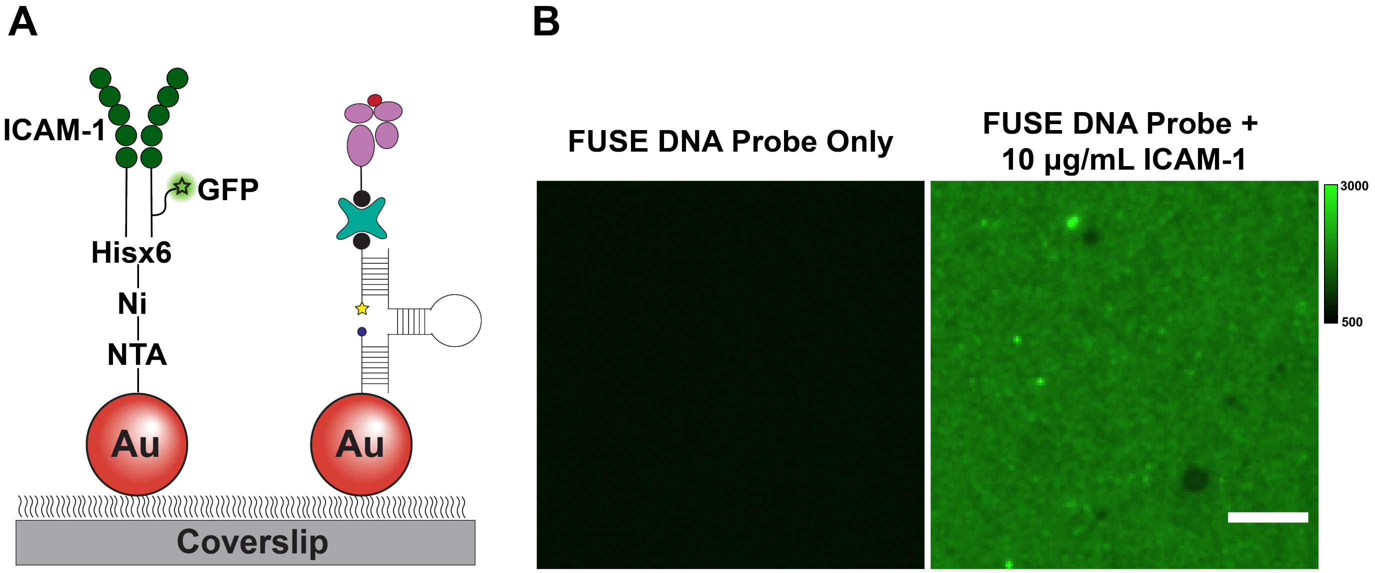

**Figure S9. GFP-ICAM-1 presentation on gold nanoparticle surface.** A) Schematic showing the surface chemistry used to present ICAM-1 alongside FUSE DNA probes on gold nanoparticle surfaces. Briefly, His-tagged ICAM-1 expressing GFP is conjugated through the imidazole-Ni-NTA interaction (see **Materials and Methods**). B) Representative images showing evidence of ICAM-1 presentation on our surfaces using GFP signal.

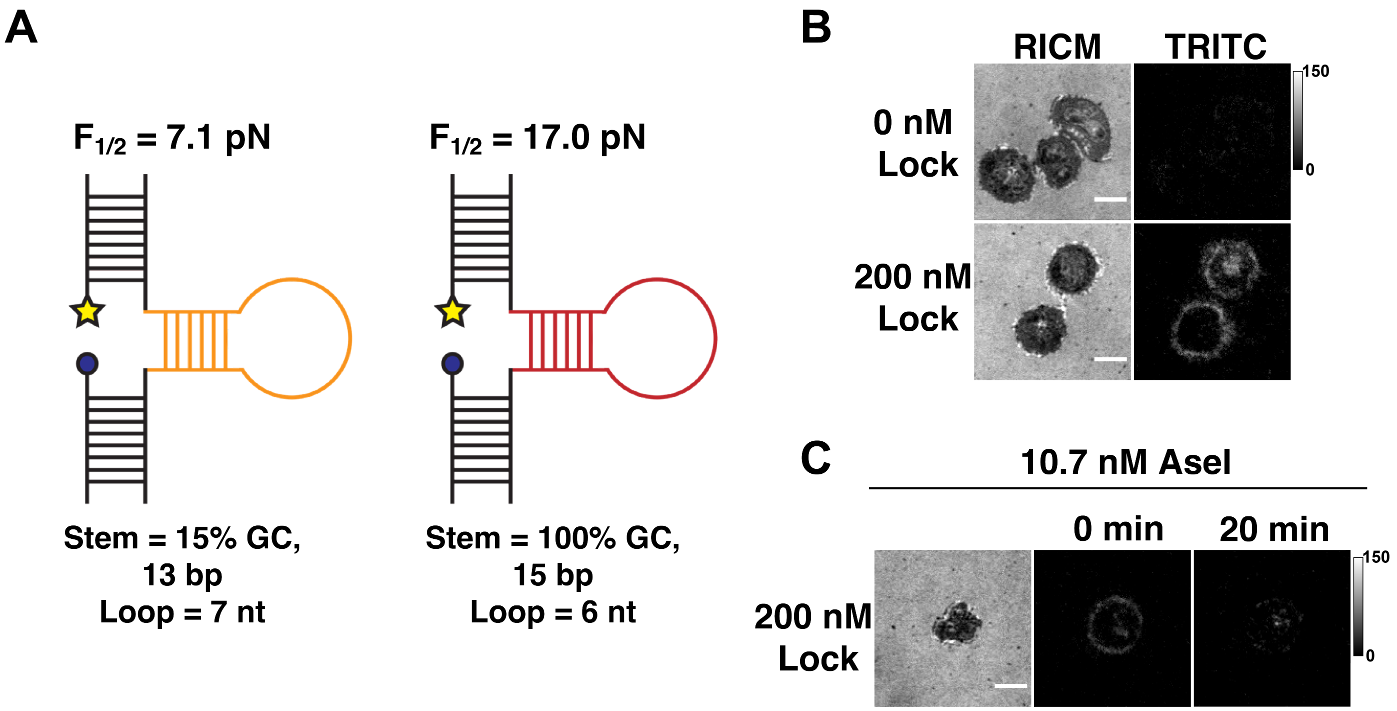

**Figure S10. Validation of 17 pN FUSE DNA probe.** A) Schematic showing the differences between the 7.1 pN and 17 pN FUSE DNA probes. The $F_{1/2}$ is increased by increasing the length and GC content of the stem, as well as decreasing the length of the loop.[1] B) Representative images showing cells spreading (RICM) and tension signal accumulating on surfaces presenting 17 pN FUSE DNA probes after 10 min incubation with or without 22mer locking strand. C) Representative images showing enzymatic cleavage of mechanically unfolded and locked 17 pN FUSE DNA probes underneath a cell after a 20 min incubation with 10.7 nM AseI.

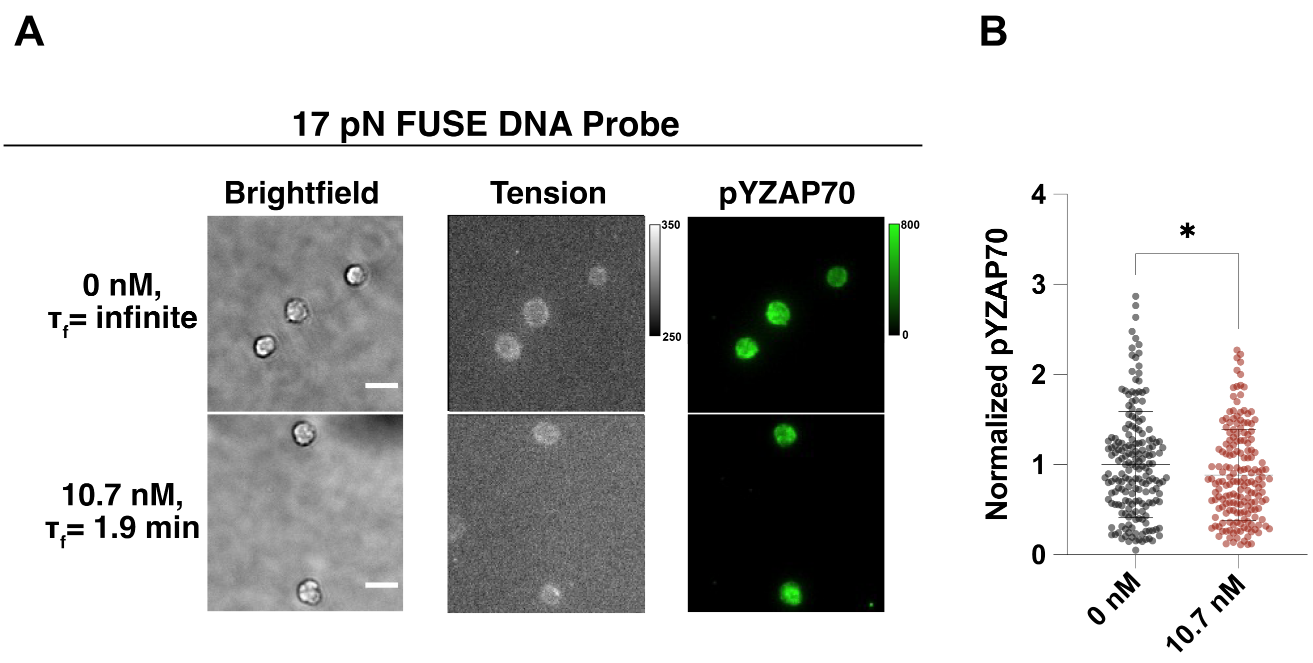

**Figure S11. Limiting $\tau_f$ of TCR-pMHC interactions with 17 pN FUSE DNA probes.** A) Representative images of cells interacting with surfaces (brightfield), tension signal, and pYZAP70 signal after 15 min incubation with or without 10.7 nM AseI on surfaces presenting N4 pMHC on 17 pN FUSE DNA probes. B) Comparison between the normalized pYZAP70 signal between the 10.7 nM (mean norm. pY = 0.88) and no enzyme conditions (mean norm. pY = 1.00), n>180 cells. Statistical analysis was performed using Student's t test, *P<0.05.
